# Systematic analysis of cotranscriptional RNA folding using transcription elongation complex display

**DOI:** 10.1101/2023.12.22.573115

**Authors:** Skyler L. Kelly, Eric J. Strobel

## Abstract

RNA can fold into structures that mediate diverse cellular functions. Understanding how RNA primary sequence directs the formation of functional structures requires methods that can comprehensively assess how changes in an RNA sequence affect its structure and function. Here we have developed a platform for performing high-throughput cotranscriptional RNA biochemical assays, called <u>T</u>ranscription <u>E</u>longation <u>C</u>omplex display (TECdisplay). TECdisplay measures RNA function by fractionating a TEC library based on the activity of cotranscriptionally displayed nascent RNA. In this way, RNA function is measured as the distribution of template DNA molecules between fractions of the transcription reaction. This approach circumvents typical RNA sequencing library preparation steps that can cause technical bias. We used TECdisplay to characterize the transcription antitermination activity of 32,768 variants of the *Clostridium beijerinckii pfl* ZTP riboswitch designed to perturb steps within its cotranscriptional folding pathway. Our findings establish TECdisplay as an accessible platform for high-throughput RNA biochemical assays.

## Introduction

RNA folding is dynamic and begins during transcription^1–3^. Despite the biological importance of structured RNAs, our understanding of how RNA folds into functional structures is limited because the interplay between RNA sequence, structure, and function is complex. Linking RNA sequence, folding, and function therefore requires methods that can assess the structural and functional consequences of thousands of RNA sequence perturbations simultaneously^4,5^. The state-of-the-art approach for high-throughput cotranscriptional RNA biochemistry couples high-throughput DNA sequencing with *in situ* transcription on an Illumina sequencer flow-cell so that fluorescence-based RNA assays can be performed for tens of thousands of RNA sequence variants in parallel^4,5^. This strategy was pioneered by the <u>Hi</u>gh-<u>T</u>hroughput <u>S</u>equencing-<u>R</u>NA <u>A</u>ffinity <u>P</u>rofiling (HiTS-RAP)^6^ and Quantitative Analysis of <u>RNA</u> on a <u>Ma</u>ssively <u>P</u>arallel Array (RNA-MaP)^7^ methods, which were initially developed to quantify RNA-protein interactions and have been collectively referred to as <u>RNA</u> array on a <u>Hi</u>gh-<u>T</u>hroughput <u>S</u>equencer (RNA-HiTS) methods. RNA-MaP assays for diverse RNA-mediated functions have since been developed^8–18^.

The primary limitation of RNA-HiTS experiments is that they require custom instrumentation. Although detailed procedures are available for building the requisite instrument^8^, many laboratories lack the expertise needed to assemble and maintain a custom instrument. To address this, we have developed a modular platform for cotranscriptional RNA biochemical assays, called <u>T</u>ranscription <u>E</u>longation <u>C</u>omplex display (TECdisplay) that quantifies the function of at least tens of thousands of RNA sequence variants simultaneously using an RNA-dependent DNA fractionation strategy that can be performed at the laboratory bench (Fig. 1a). In a TECdisplay experiment, an RNA variant library is cotranscriptionally displayed from *E. coli* RNA polymerase (RNAP) using quantitative *in vitro* transcription. This TEC library is then fractionated by the function of the nascent RNA. Because template DNA is physically coupled to nascent RNA in the TEC, the distribution of template DNA molecules for a given variant between fractions reflects the function of the RNA that it encodes. Consequently, template DNA can be recovered, quantitatively tagged with fraction-specific barcodes, and sequenced to determine the activity of each RNA variant in the library.

**Figure 1.**
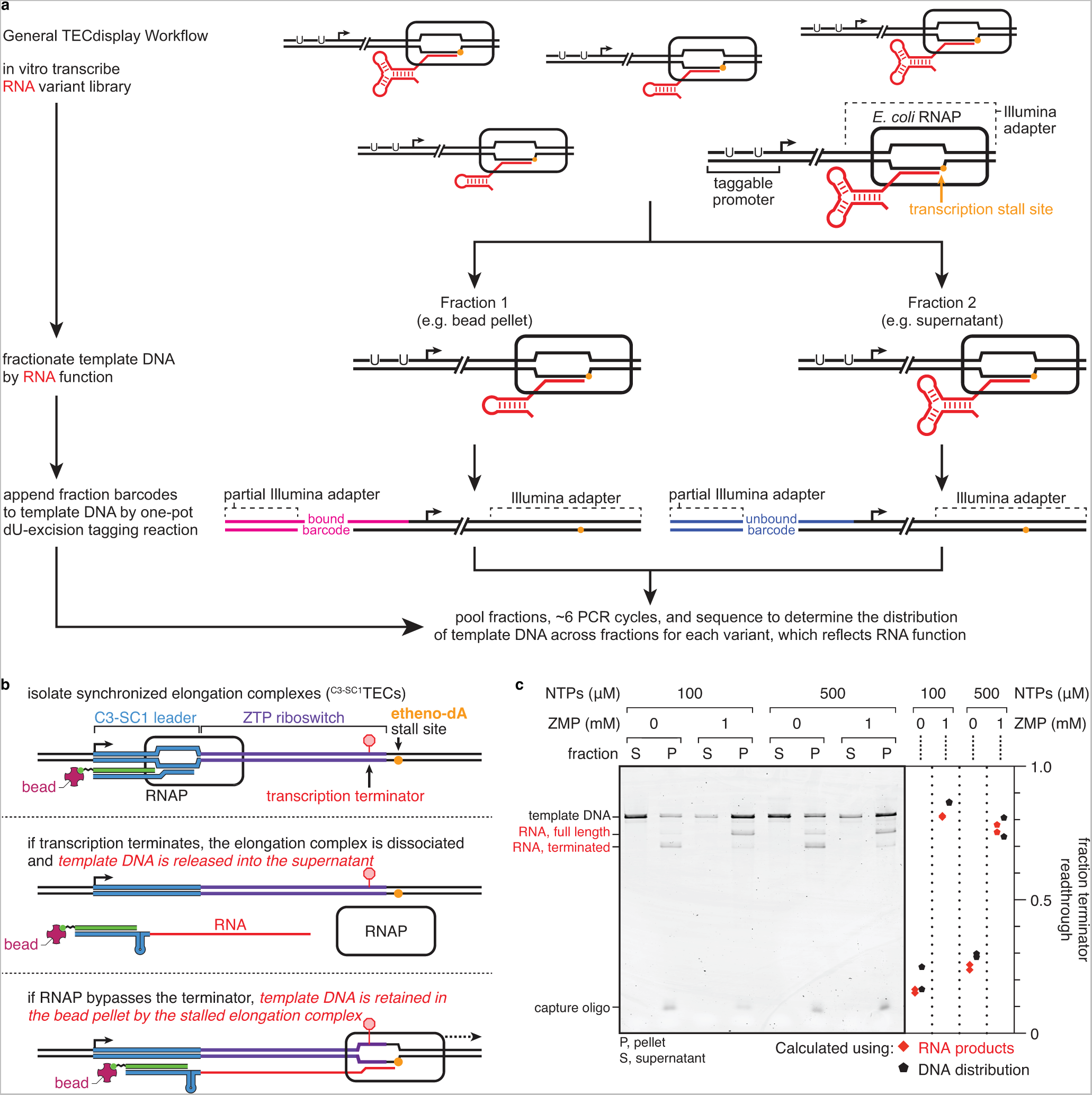
Overview of the TECdisplay procedure. **a**, General TECdisplay workflow. A library of RNA variants is encoded in template DNA that contains a promoter that is compatible with dU-excision tagging and a transcription stall site that prevents RNAP from running off the template. Single-round transcription is performed under conditions that yield TECs with a 1:1:1 DNA:RNAP:RNA composition. The TEC library is then fractionated by RNA function. This fractionation step can vary depending on the RNA function being measured. The protein and RNA components of the transcription reaction are then degraded and fraction-specific barcodes are quantitatively appended to template DNA using the one-pot dU-excision tagging procedure. Barcoded fractions are pooled together, amplified for ∼6 cycles of PCR, and sequenced to determine the distribution of template DNA across fractions for each variant in the library. Because DNA was fractionated by RNA function, the distribution of DNA between fractions reflects RNA function. **b**, Strategy for measuring riboswitch-mediated transcription termination/antitermination activity using TECdisplay. TECs are synchronized using the C3-SC1 leader sequence and immobilized on streptavidin-coated magnetic beads by a biotinylated capture oligonucleotide. An etheno-dA transcription stall site is positioned downstream of the riboswitch intrinsic terminator to retain RNAP on DNA. When transcription is resumed, template DNA molecules on which a termination event happened are released into the supernatant. Conversely, template DNA molecules on which a terminator readthrough event happened are retained in the bead pellet. **c**, Quantification of *C*. *beijerinckii pfl* ZTP riboswitch activity in various conditions using the fractionation strategy shown in panel b. Fraction terminator readthrough was calculated using the distribution of template DNA between supernatant and pellet fractions and using the distribution of terminated and full length RNA for each sample. The gel image is representative of n=2 replicates. RNAP, RNA polymerase.

Here we describe the first TECdisplay assay, which measures riboswitch-mediated transcription termination efficiency. Riboswitches are cis-acting RNA chemical sensors that regulate gene expression in response to diverse ligands^19,20^ and have been used to develop biomolecular sensors^21,22^ and chemical biology tools^23,24^. We used TECdisplay to quantify the transcription termination and antitermination activity of 32,768 variants of the *Clostridium beijerinckii* (*C. beijerinckii*, Cbe) *pfl* ZTP riboswitch designed to perturb ZTP aptamer and transcription terminator folding. The ZTP riboswitch, which regulates genes associated with purine biosynthesis in response to the metabolites ZTP and ZMP^25^, is a potential antibiotic target^26^ and has been used as a reporter in a screen for folate biosynthesis inhibitors^27^. Our analysis of ZTP riboswitch folding produced expected outcomes that validate the accuracy of TECdisplay, identified sequence combinations that would naively be predicted to be functional based on sequence but are non-functional due to misfolding, and uncovered how stabilization of a non-canonical base pair increases ZMP-responsiveness. Our findings establish TECdisplay as an accessible platform for high-throughput cotranscriptional RNA functional assays that can likely be used to characterize diverse RNA-mediated biochemical activities.

## Results

### The cotranscriptional RNA-dependent DNA fractionation strategy

TECdisplay is a modular platform for performing high-throughput cotranscriptional RNA biochemical assays. TECdisplay experiments comprise three steps: First, an RNA variant library is synthesized using a quantitative *E. coli* single-round *in vitro* transcription reaction in which virtually all DNA template molecules yield a single RNA product^28^ (Fig. 1a). This is accomplished using the C3-SC1 leader sequence to purify synchronized ^C3-SC1^TECs, which are >98% active^28^. The template DNA used for this reaction invariably contains one of several modifications that halt *E. coli* RNAP transcription, which prevents run-off transcription so that nascent RNA molecules are physically coupled to the DNA template from which they were transcribed^29^. Second, the TEC library is fractionated by the biochemical activity of the nascent RNA (Fig. 1a). For example, in this work we describe a TECdisplay assay for measuring transcription termination efficiency. In this assay, RNAP is tethered to a magnetic bead by the Cap3 oligonucleotide, which anneals to the C3 hybridization site in the C3-SC1 leader transcript^28^, and an etheno-dA transcription stall site^29,30^ is positioned downstream of the transcription termination site (Fig 1b). If transcription termination occurs, nascent RNA is released from the TEC and template DNA is released into the supernatant. Conversely, if RNAP bypasses the termination site, transcription halts at the etheno-dA stall site and the template DNA is retained in the bead pellet. In this way, DNA molecules are separated based on whether transcription termination occurred so that the efficiency of terminator readthrough can be calculated using the distribution of template DNA across supernatant and pellet fractions (Fig. 1c). This fractionation step is modular and can be changed depending on the RNA biochemical activity that will be measured. Third, the template DNA is converted into an Illumina sequencing library. In this step, RNA and protein are degraded and fraction- and molecule-specific barcodes are quantitatively appended to the template DNA in a one-pot <u>d</u>eoxy<u>U</u>ridine e<u>X</u>cision tagging (dUX-tagging) reaction^31^ (Fig. 1a, Supplementary Fig. 1a). Following the dUX-tagging reaction, the pellet and supernatant fractions are pooled together and excess tagging oligonucleotide is depleted by solid-phase reversible immobilization (SPRI). The non-transcribed DNA strand is then degraded using lambda exonuclease so that only the transcribed DNA strand, which was used to generate the nascent RNA, will be sequenced (Supplementary Fig. 1). This step safeguards the experiment against the potential presence of heteroduplexes in the DNA template preparation, which would confound the experimental measurement. The barcoded template DNA is then amplified by ∼6 cycles of PCR. The resulting library is then sequenced using the Illumina platform to determine the distribution of template DNA between pellet and supernatant fractions for each variant, which reflects the functional properties of the nascent RNA.

The RNA-dependent DNA fractionation strategy used by TECdisplay is advantageous in two ways. First, TECdisplay circumvents typical RNA sequencing library preparation steps like reverse transcription and single-stranded nucleic acid ligations, which can potentially introduce technical bias into the measurement^32–39^. Second, TECdisplay is compatible with destructive RNA functions in which RNA sequence information is lost during the assay because sequence information is recovered directly from the DNA.

### Systematic perturbation of *pfl* ZTP aptamer folding

To validate TECdisplay, we characterized the activity of 32,768 *Clostridium beijerinckii pfl* ZTP riboswitch variants in which conserved purine and pyrimidine nucleotides within the ZTP aptamer P1 stem, pseudoknot (PK), and terminator and are randomized (Fig. 2a). This library perturbs the competition between terminator and pseudoknot folding, which has several predictable outcomes and was previously assessed in a mid-throughput analysis of 64 *Cbe pfl* ZTP riboswitch variants^40^. The current data set, which contains 512 times more variants, enabled us to confidently dissect how sequence variation in pseudoknot and terminator base pairs dictates RNA folding outcome because the same competition between pseudoknot and terminator folding was observed repeatedly for different P1 stem variants. Comparison of replicates collected with 0 mM (R^2^ = 0.987) or 1 mM ZMP (R^2^ = 0.989) shows that TECdisplay measurements are reproducible (Fig. 2b). Bulk visualization of ZMP-mediated antitermination efficiency in response to increasing ZMP concentration revealed that the variant library contains discrete clusters of non-functional variants with variable terminator efficiency and functional variants with broadly variable terminator efficiency and responsiveness to ZMP (Fig. 2c). Filtering the 0 mM and 1 mM ZMP data for variants in which P1, PK, and terminator base pairs are intact but not constrained by identity enriches for functional variants, although many non-functional variants remain (Fig. 2d, e). Filtering these data for variants in which variable nucleotides in PK and the terminator form Watson-Crick pairs isolates a set of functional riboswitches in which the increase in fraction terminator readthrough in the presence of 1 mM ZMP is at least 0.2 for 94.5% of the variants (Fig. 2e, f). In agreement with this observation, the ZMP dose response for select variants matches four expectations: First, variants with intact PK and terminator base pairs exhibit ZMP-mediated transcription antitermination (Fig. 2g). Second, variants with a disrupted PK and intact terminator terminate transcription efficiently but do not respond to ZMP (Fig. 2h). Third, variants with an intact PK and disrupted terminator fail to terminate transcription efficiently regardless of ZMP concentration (Fig. 2i). Fourth, variants in which both PK and the terminator are disrupted do not respond to ZMP and terminate transcription more efficiently than when only the terminator is disrupted, which agrees with a previous finding that PK is a barrier to terminator base pair propagation (Fig. 2j). Furthermore, the discrete clusters of non-functional variants are attributable to distinct classes of variants defined by the identity of pseudoknot and terminator base pairs (Supplementary Fig. 2) Together, these data indicate that the TECdisplay assay for transcription termination efficiency accurately measures riboswitch activity.

**Figure 2.**
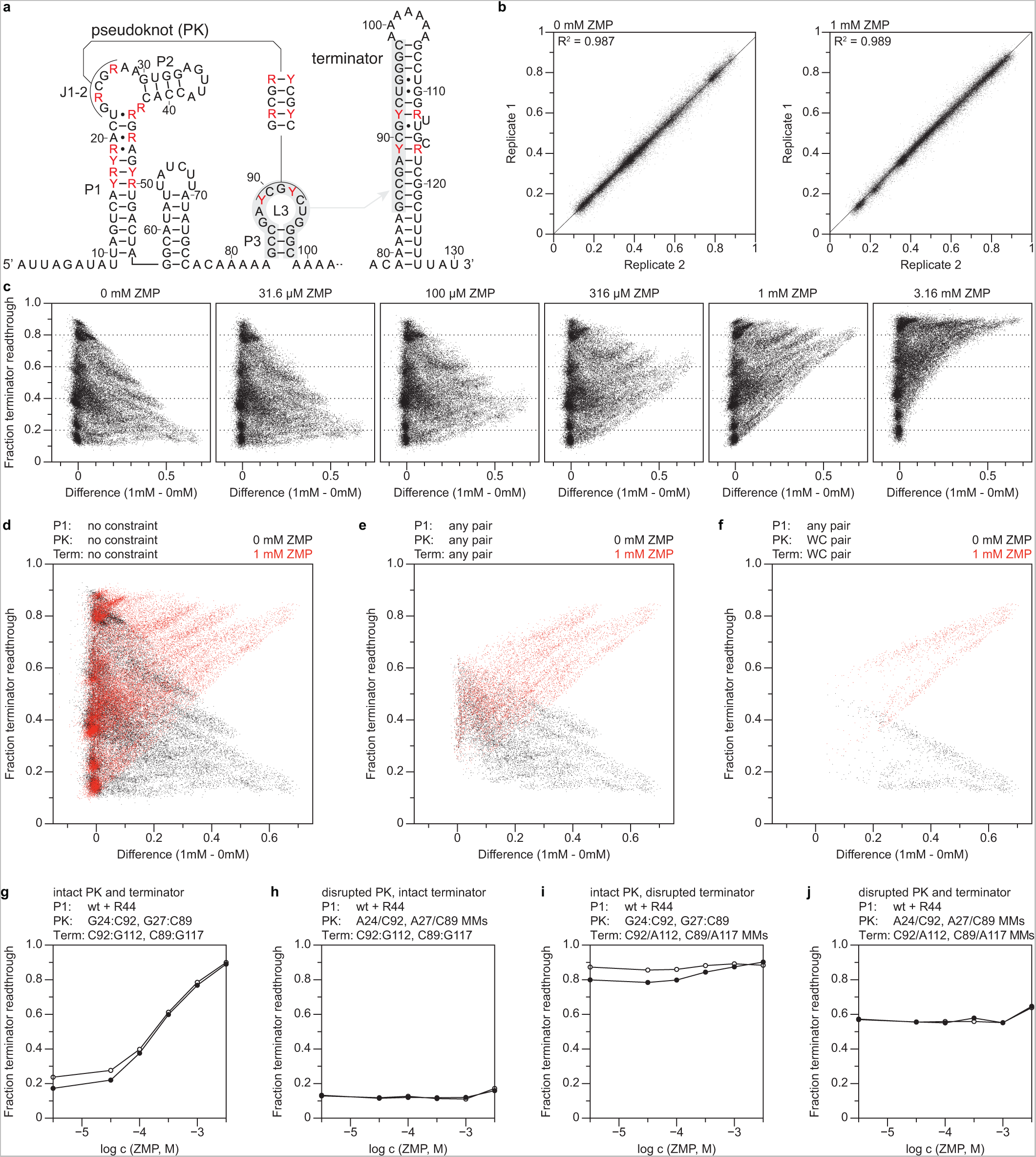
Overview of the *C. beijerinckii pfl* ZTP riboswitch consensus variant library. **a**, Sequence of the *C*. *beijerinckii pfl* ZTP riboswitch consensus variant library. **b**, Correlation of replicate TECdisplay fraction terminator readthrough measurements for the 0 mM and 1 mM ZMP conditions. **c**, Plots showing fraction terminator readthrough for the sequence variants shown in panel a as measured by TECdisplay with variable ZMP concentration. Variants are distributed across the x-axis by the difference in fraction terminator readthrough measured for the 1 mM ZMP and 0 mM ZMP conditions so that each variant is located at the same x-axis position in all plots. **d-f**, Plots showing fraction terminator readthrough for the 0 mM ZMP (black) and 1 mM ZMP (red) conditions for all variants (**d**), variants in which P1, PK, and the terminator are fully paired without base pair identity constraints (**e**), and variants in which P1, PK, and the terminator are fully paired but PK and the terminator contain only Watson-Crick base pairs (**f**). **g-j**, Plots showing fraction terminator readthrough as a function of ZMP concentration for wt-like variants with strong pseudoknot and terminator pairs (**g**), pseudoknot mismatches and strong terminator pairs (**h**), strong pseudoknot pairs and terminator mismatches (**i**), and pseudoknot and terminator mismatches (**j**). TECdisplay data for the 0 mM and 1 mM ZMP conditions are from n=2 replicates that were combined. TECdisplay data for the 31.6 μM, 100 μM, 316 μM, and 3.16 mM conditions are n=1. PK, pseudoknot; term, terminator; WC, Watson-Crick; MM, mismatch.

### Differential terminator activity of *pfl* riboswitch pseudoknot variants is caused by misfolding

Filtering the 0 mM and 1 mM *pfl* riboswitch data sets for variants in which PK and the terminator comprise Watson-Crick base pairs and P1 is intact isolated a set of variants that appears to contain at least two subpopulations (Fig. 2f). The *pfl* riboswitch consensus variant library contains four possible PK/terminator configurations, which are referred to as wild-type (WT), flip (FL), strong (ST), and weak (WK) in the text below (Fig 3a). Sub-elements of these classes are referred to as ‘sub-element_class_’. For example, the wild-type J1-2 and terminator sub-elements are referred to as J1-2_WT_ and Term_WT_, respectively. Filtering the variants identified in Figure 2f by these classes revealed that each PK/terminator configuration forms a distinct cluster, but that there is a relationship between the terminator efficiency of WT and WK variants and between FL and ST variants (Fig. 3b). 97% of WT and WK variants exhibit terminator efficiencies between ∼75% and ∼90%, and the efficiency of termination is independent of the ZMP-mediated antitermination response (Fig. 3b). In contrast, FL and ST variants exhibit terminator efficiencies ranging from ∼50% to ∼90% and the amplitude of the ZMP-mediated antitermination response correlates with terminator efficiency (Fig. 3b). Whereas FL and ST variants contain non-native G24:C92 and C92:G112 base pairs, WT and WK variants contain the native A24:U92 and U92:A112 base pairs (Fig 3a). To identify how these nucleotides contribute to the termination defects observed for FL and ST variants, we assessed how each possible J1-2 and terminator hairpin combination affected terminator efficiency in the absence of ZMP (Fig 3c). In the presence of J1-2_WT_ (A24, G27) or J1-2_WK_ (A24, A27), Term_FL_ and Term_ST_ terminate transcription comparably to the cognate terminators (Fig. 3c, groups 1-3 and 14-16). This shows that neither Term_FL_ nor Term_ST_ is intrinsically defective. This indicates that the identity of position 24, which distinguishes J1-2_WT_ and J1-2_WK_ (A24) from J1-2_FL_ and J1-2_ST_ (G24), contributes to the termination defect observed for FL and ST riboswitches while the identity of the 92:112 base pair does not.

**Figure 3.**
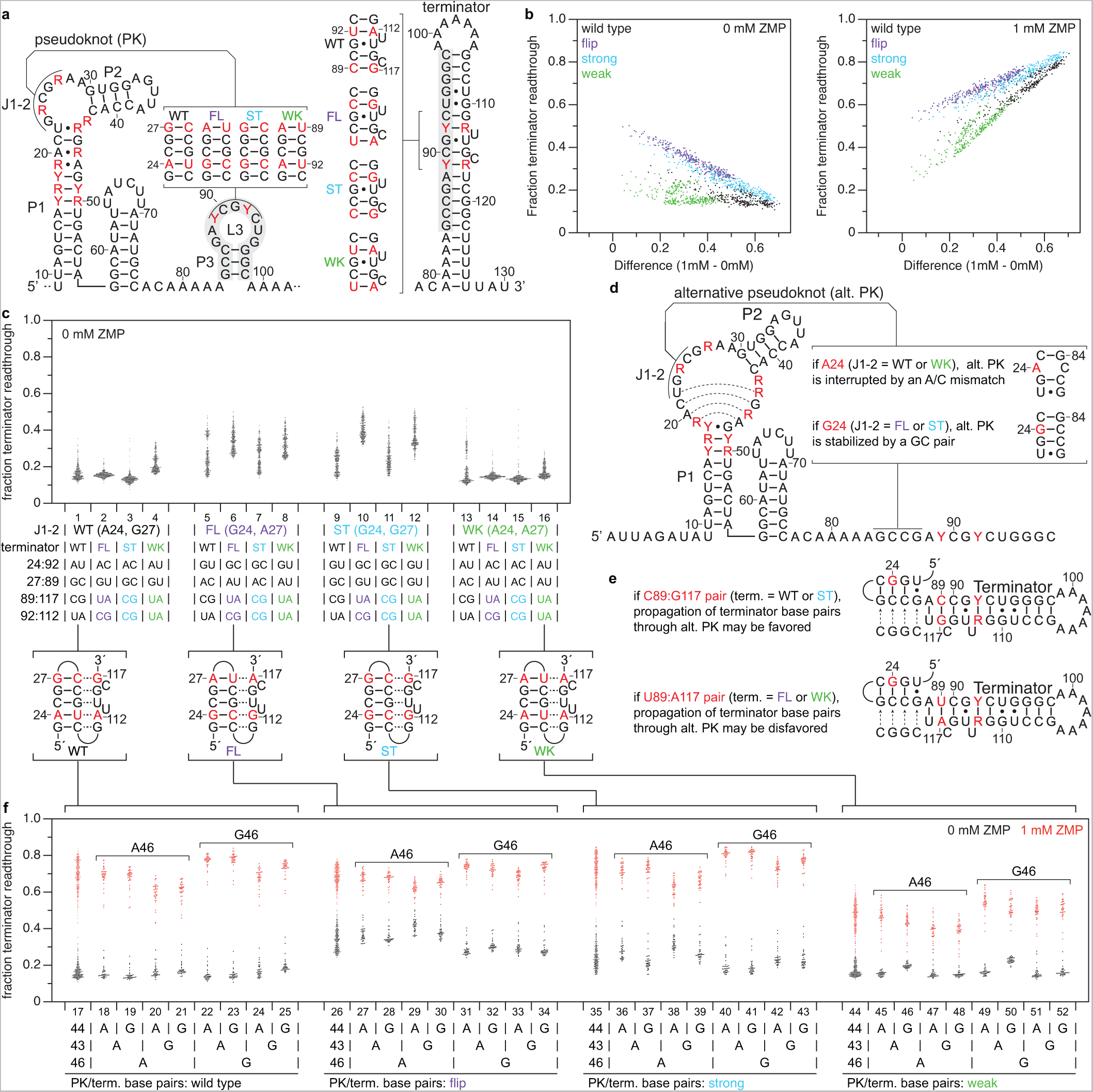
Activity of *C. beijerinckii pfl* ZTP riboswitch intact pseudoknot and terminator variants. **a**, Sequence of the *C*. *beijerinckii pfl* ZTP riboswitch consensus variant library annotated to show all pseudoknot and terminator configurations in which variable nucleotides form Watson-Crick base pairs. **b**, Plots showing fraction terminator readthrough for the four variant classes in which variable nucleotides in the pseudoknot and terminator form Watson-Crick base pairs. Variants are distributed across the x-axis by the difference in fraction terminator readthrough measured for the 1 mM ZMP and 0 mM ZMP conditions so that each variant is located at the same x-axis position in both plots. **c**, Plot showing the efficiency of each terminator from panel a in the presence of each J1-2 sequence. **d**, Illustration of an alternative pseudoknot that is favored if J1-2 contains G24. **e**, Illustration of a model for how the identity of the 89:117 terminator base pair could affect propagation of terminator base pairs through the alternative pseudoknot depicted in panel d. **f**, Plot showing how the composition of nucleotides 43, 44, and 46 affect fraction terminator readthrough for the four variant classes in which variable nucleotides in the pseudoknot and terminator form Watson-Crick base pairs. All data are from the experiment shown in Figure 2. WT, wild type; FL, flip; ST, strong; WK, weak; PK, pseudoknot; term, terminator.

The distribution of terminator efficiencies was broader when J1-2 contained G24 regardless of which terminator hairpin was present (Fig. 3c, groups 5-12). Nonetheless, two populations emerged: variants that contain G24 terminate transcription more efficiently in the presence of Term_WT_ or Term_ST_ than in the presence of Term_FL_ or Term_WK_ (Fig. 3c, groups 5-12). This indicates that the identity of the 89:117 pair, which distinguishes Term_WT_ and Term_ST_ (C89:G117) from Term_FL_ and Term_WK_ (U89:A117), also contributes to the termination deficiencies observed for FL and ST riboswitches.

The observation that G24 inhibits both transcription termination and ZMP-mediated antitermination suggests that it can cause the *pfl* aptamer to misfold. One possible mechanism for misfolding is the formation of an alternative pseudoknot between nucleotides 22-25 and 87-84, which is stabilized by G24 and interrupted by A24 and is adjacent to the native pseudoknot (Fig. 3d). This model is consistent with the observation that a U89:A117 terminator base pair exacerbates the termination defect caused by G24. Displacement of the alternative pseudoknot during terminator base pair propagation is anchored by two contiguous base pairs due to a bulge at C116 (Fig. 3e). The presence of a strong base pair within the anchoring dinucleotide may favor displacement of the alternative pseudoknot.

In addition to the variability in terminator activity between the four PK/terminator classes, variants within each class exhibit a variable ZMP-mediated antitermination response (Fig. 3b). This variability is primarily attributable to the identity of nucleotides 43, 44, and 46, which are present in a region of P1 that contains several non-canonical base pairs (Fig. 3a). In variants in which variable PK and terminator nucleotides form Watson-Crick pairs, variants that contain G46 and A43 exhibit a stronger antitermination response (Fig. 3f). The identity of position 44 is associated with context-dependent effects on transcription termination efficiency in variants that combine non-cognate J1-2 and terminator configurations (Supplementary Fig. 3). Before investigating the contribution of nucleotides 46 and 43 to ZMP-mediated transcription antitermination, it was necessary to account for a subset of variants in which the ZTP aptamer is prone to misfolding due to the presence of competitor helices, which is described in the section below.

### Competitor helices can cause aptamer misfolding

Most variant groups identified in Figure 3f exhibit a fraction terminator readthrough distribution that contains a tail comprised of variants with increased ZMP-independent terminator readthrough. Minimum free energy structure prediction^41^ revealed that variants within these tails have the capacity to form two competitor helices, termed CH1 and CH2, that are mutually exclusive with the native P1 subdomain structure (Fig. 4a, b). With the exception of WK variants that contain A43, which do not appear to misfold, filtering for variants with the capacity to form CH1 and CH2 isolates the tail of each distribution (Fig. 4c and Supplementary Fig. 4c, filter 1). Disrupting either CH1 or CH2 restores terminator efficiency, indicating that both competitor helices must be present to cause aptamer misfolding (Fig. 4c and Supplementary Fig. 4c, filters 2 and 3).This indicates that CH1/CH2 is responsible for aptamer misfolding that affects terminator efficiency. To assess whether PK formation is required for the CH1/CH2-dependent termination defect, we applied the filters shown in Figure 4b to *pfl* riboswitch variants in which PK contains at least 1 mismatch (Figure 4d, e). For most variants that contain CH1/CH2, disrupting PK eliminates the transcription termination defect to the same degree as disrupting CH1 or CH2 (Fig. 4d, e). However, in the presence of Term_ST_, variants that exhibit a CH1/CH2-dependent terminator defect in the absence of PK all contain G43 and G44, which permits the formation of an alternative pseudoknot with L3. Similarly, in the presence of Term_FL_, variants that exhibit a CH1/CH2-dependent terminator defect in the absence of PK contain G43 and R44, which permits the formation of a weaker alternative pseudoknot with L3. This trend was observed throughout all variant classes in which PK was disrupted and suggests that PK formation is required for CH1/CH2 to inhibit transcription termination, but that PK identity is not critical (Supplementary Fig. 2).

**Figure 4.**
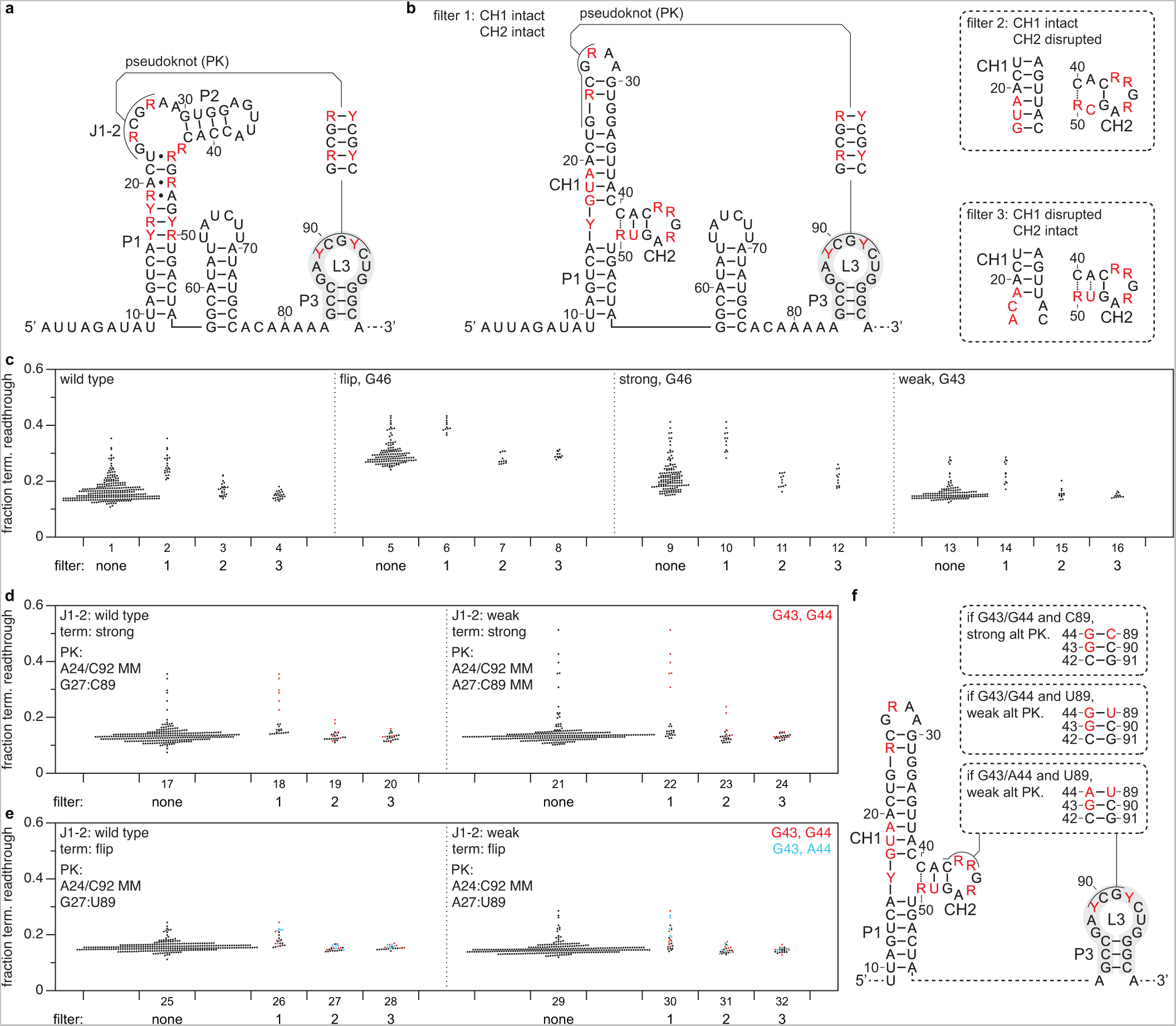
Activity of *C. beijerinckii pfl* ZTP riboswitch competitor helix variants. **a**, Sequence of the *C*. *beijerinckii pfl* ZTP riboswitch consensus variant library. **b**, Illustration of the filters that were applied in panels c-e. Filter 1 includes variants in which both CH1 and CH2 are intact. Filter 2 includes variants in which CH1 is intact and CH2 is disrupted. Filter 3 includes variants in which CH1 is disrupted and CH2 is intact. **c**, Plots showing the effect of CH1 and CH2 on fraction terminator readthrough in the absence of ZMP for variants in which variable nucleotides in the pseudoknot and terminator form Watson-Crick base pairs. Variants in groups 5-12 also contained G46 and variants in groups 13-16 also contained G43; the corresponding A46 and A43 variants are shown in Supplementary Fig. 4. **d, e**, Plots showing the effect of CH1 and CH2 on variants that contain pseudoknot mismatches. Variants with the capacity to form alternative pseudoknots are colored red and blue. **f**, Illustration of alternative pseudoknots that may form between the CH2 loop and L3. All data are from the experiment shown in Figure 2. PK, pseudoknot; term, terminator; CH1, competitor helix 1; CH2, competitor helix 2; MM, mismatch.

### P1 base pairs modulate ZMP-mediated transcription antitermination

Isolating the effect of CH1/CH2 enabled the investigation of how nucleotides 46 and 43 contribute to the ZMP-mediated antitermination response. To assess how nucleotides 46 and 43 affect transcription antitermination, we filtered WT PK variants for each of the four possible sequence configurations of positions 46 and 43 and generated aggregate dose-response curves (Fig. 5a). As expected, the presence of the CH1/CH2 competitor helix increases ZMP-independent terminator readthrough and inhibits ZMP-mediated antitermination in all cases (Fig. 5a). Comparison of aggregate response curves indicates a sequence preference of [G46, A43] > [G46, G43] ≈ [A46, A43] > [A46, G43] (Fig. 5b). The preference for G46, which is non-native and pairs with A20 in P1, is presumably attributable to P1 stabilization, since G46 forms three hydrogen bonds with A20, whereas A46 only forms two hydrogen bonds with A20 (Fig. 5c). The preference for A43, which is native and stacks with the U22: R44 base pair, is less clear. The structure of the *Actinomyces odontolyticus* ZTP aptamer determined by Trausch et al. identified a base pair between the P2-proximal adenine base in J1-2 and a guanine base at the position equivalent to nucleotide 43 in the *Cbe pfl* ZTP aptamer^42^ (Fig. 5d). In the *Cbe pfl* aptamer, the DMS reactivity of the P2-proximal adenine at position 29 increases upon ZMP binding^38^ (Fig. 5e). This indicates that ZMP binding causes the *Cbe pfl* aptamer to undergo a conformational change in which the Watson-Crick face of A29 becomes unpaired. An interaction between A29 and G43 might interfere with the ZMP-mediated conformational change that causes A29 to become DMS reactive. However, this speculation cannot be assessed rigorously by TECdisplay or cotranscriptional RNA chemical probing. Nonetheless, it is clear that the nucleotide composition of P1 can modulate the ZMP-mediated transcription antitermination response.

**Figure 5.**
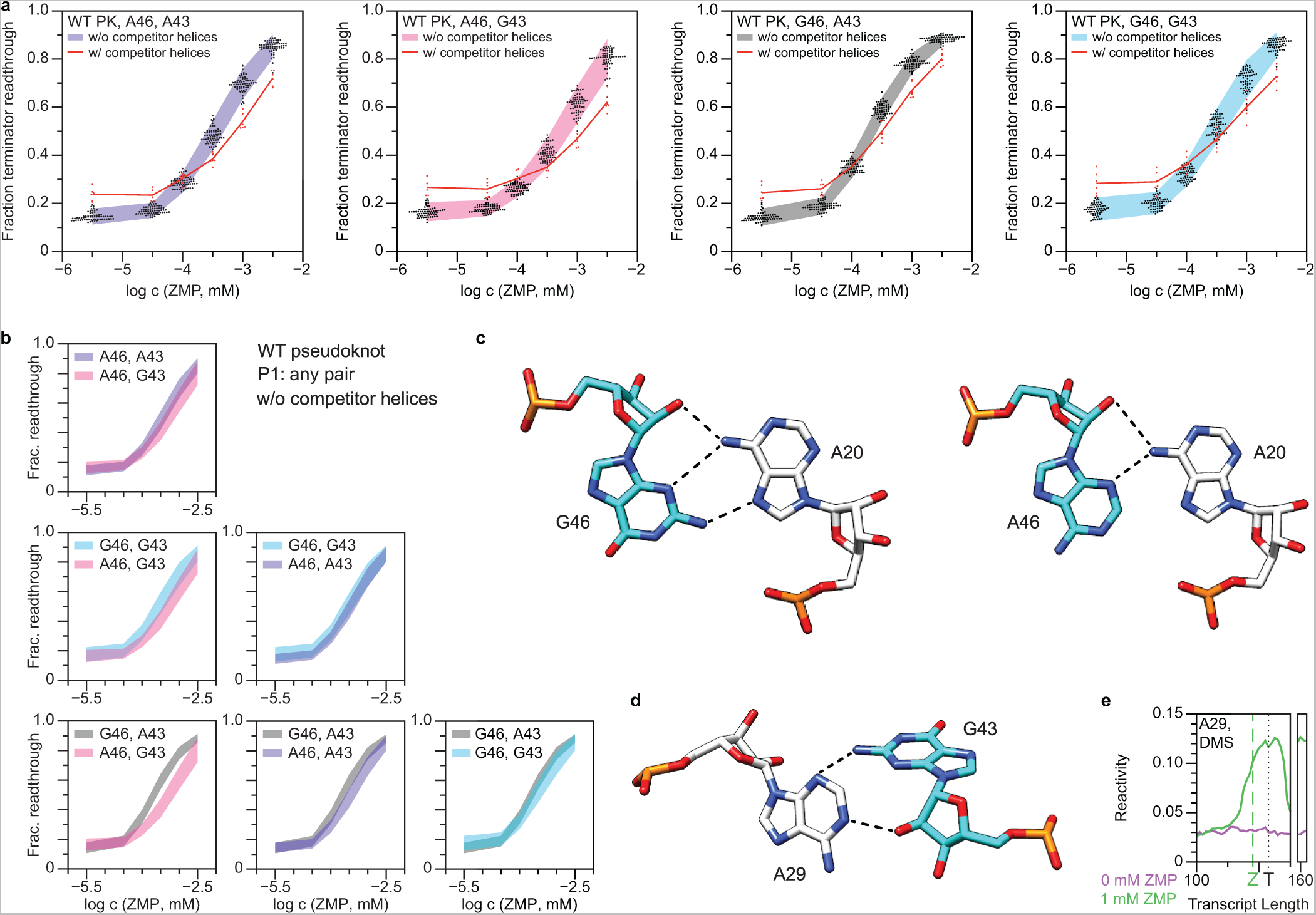
Contribution of P1 nucleotides to ZMP-responsiveness. **a,** Plots showing the ZMP-mediated transcription antitermination response for *C. beijerinckii pfl* ZTP riboswitch variants with intact P1 base pairs and wild-type pseudoknot and terminator base pairs filtered by the identity of nucleotides 46 and 43. Variants without competitor helices (black) are shown with a shaded region indicating the mean fraction terminator readthrough ± two standard deviations. Variants with competitor helices are shown in red. Data are from the experiment in Figure 2. **b,** Plots comparing the ZMP-mediated antitermination response of the four classes of variants shown in panel a. **c,** Structure of the *Thermosinus carboxydivorans* (PDB:4ZNP^69^) and *Fusobacterium ulcerans* (PDB:5BTP^70^) ZTP aptamer base pairs that correspond to the *Cbe* A20:G46 and A20:A46 base pairs, respectively. **d,** Structure of the *Actinomyces odontolyticus* ZTP aptamer (PDB:4XW7^42^) base pair that corresponds to the *Cbe* A29:G43 base pair. Crystal structures were visualized using UCSF Chimera^71^ and are labeled using *Cbe* numbering. **e,** Plot of transcript length-dependent DMS reactivity changes at A29 measured using TECprobe-VL. Z indicates when ZMP can bind to the aptamer and T indicates when the terminator hairpin can begin to fold. Decreased A29 DMS reactivity at transcript ∼130 is caused by the presence of terminated transcripts in the probing reaction. TECprobe-VL data are from Szyjka and Strobel, 2023, *Observation of coordinated RNA folding events by systematic cotranscriptional RNA structure probing*, *Nat. Commun.*, 2023 Nov 29; 14(1)7839^38^. WT, wild type; frac., fraction.

## Discussion

We have described how TECdisplay quantifies riboswitch-mediated transcription antitermination using a cotranscriptional RNA-dependent DNA fractionation strategy. While the current study focuses on the measurement of transcription termination efficiency, TECdisplay is a modular platform for building assays that measure diverse RNA-mediated functions. As outlined in Figure 1, the general TECdisplay procedure records RNA function in an Illumina sequencing library in three steps: First, RNA is cotranscriptionally displayed from *E. coli* RNAP using a quantitative single-round *in vitro* transcription reaction in which RNAP is halted at a chemically-encoded transcription roadblock. Second, TECs are fractionated by the activity of the cotranscriptionally displayed transcript. Third, template DNA molecules are quantitatively tagged with fraction-specific barcodes. The fractions are then pooled back together and amplified for Illumina sequencing by limited-cycle PCR. This modular organization is designed to facilitate the rapid development of new TECdisplay assays. Whereas the procedures for quantitatively barcoding template DNA and amplifying an Illumina library are universal elements of all TECdisplay assays, the procedures for cotranscriptionally displaying RNA and fractionating the TEC library are designed to be tailorable to the application at hand. Towards this end, we have already established a high-performance tool kit for building cotranscriptional RNA assays. While the current procedure uses an etheno-dA modification to halt RNAP, we previously showed that the biotin-triethylene glycol and desthiobiotin-triethylene glycol affinity tags quantitatively halt *E. coli* RNAP transcription when placed in the transcribed DNA strand^29^. Similarly, Nadon et al. showed that the NPOM-dT modification can be used as a reversible transcription roadblock^43^. The biotin-triethylene glycol modification is likely to be particularly useful for TECdisplay because it can be used to isolate roadblocked TECs with a 1:1:1 DNA:RNAP:RNA composition without using the non-native C3-SC1 leader sequence^44,45^. We envision that these tools will facilitate the development of TECdisplay assays for diverse RNA functions.

TECdisplay is directly inspired by the HiTS-RAP^6^ and RNA-MaP^7^ (RNA-HiTS) methods, in which RNA is cotranscriptionally displayed *in situ* on an Illumina flow cell for use in high-throughput, fluorescence-based RNA functional assays. Like HiTS-RAP and RNA-MaP, TECdisplay enables the activity of at least tens of thousands of RNA variants to be measured simultaneously. The primary advantage of TECdisplay is that it is broadly accessible because it does not require custom instrumentation and can be performed using commercially available reagents with the exception of GreB, which is readily purified^46^. The primary disadvantage of TECdisplay relative to RNA-HiTS methods is that TECdisplay is not a direct biophysical measurement. Consequently, RNA-HiTS experiments have superior temporal-resolution and may be more quantitatively accurate in some cases. While there is substantial overlap in the potential applications of TECdisplay and RNA-HiTS assays, these strategies are also likely to be complementary due to the technical nuances of implementing fractionation-based and imaging-based assays for RNA function.

Torgerson et al., previously developed a high-throughput transcription termination assay called <u>S</u>equencing-based <u>M</u>utational <u>A</u>nalysis of <u>R</u>NA <u>T</u>ranscription <u>T</u>ermination (SMARTT)^47^, which has been used to characterize mechanisms of riboswitch-mediated transcription control^48^ and was integrated with a RelA cleavage assay^49^ to measure riboswitch-mediated translation control^50^. SMARTT quantifies transcription termination efficiency using an RNA sequencing approach in which sequencing reads that originate from termination products are distinguished from full length reads by the identity of their 3’ end. The TECdisplay assay presented here is also designed to systematically quantify transcription termination efficiency, but accomplishes this measurement using an orthogonal fractionation-based approach. TECdisplay is a more complex procedure than SMARTT because it is designed to be a modular platform for developing cotranscriptional RNA assays rather than a dedicated assay for transcription termination efficiency. For some RNA targets, the TECdisplay assay for quantifying transcription termination efficiency described here will be advantageous because it does not require an RNA 3’ ligation which can be a source of technical bias^32,33,36,38^. In cases where the target RNA cannot accommodate the C3-SC1 leader sequence, which is required for the TECdisplay assay, SMARTT will be advantageous because it does not require a non-native leader sequence.

There are several potential sources of error that must be avoided to ensure that cotranscriptional RNA-dependent DNA fractionation is robust during TECdisplay experiments. First, because sequence information for each variant is recovered from DNA, heteroduplexes in the DNA template preparation will confound TECdisplay measurements. Strategies such as performing reconditioning PCR cycles can minimize the presence of heteroduplexes^51^. To further safeguard against variability in the success of DNA template reconditioning, the non-transcribed DNA strand is degraded by lambda exonuclease after template DNA fractions are barcoded and pooled back together during the TECdisplay sequencing library preparation. This guarantees that all sequencing reads are derived from the transcribed DNA strand, which directly corresponds to the RNA sequence. Second, it is essential that virtually every DNA template molecule in the transcription reaction in the yields a single, complete RNA product because DNA templates that do not contain a single TEC that has completed transcription will partition into one reaction fraction by default and cause noise in the TECdisplay measurement. Quantitative *in vitro* transcription is accomplished by isolating ^C3-SC1^TECs^28,52^, which are synchronized and >98% active and by including the transcript cleavage factor GreB^53^, which rescues TECs that have arrested due to RNAP backtracking^54–56^, in the transcription reaction. It is also possible for long-lived pauses that are not backtracked^57–59^ to prevent RNAP from completing transcription, however this can typically be resolved by increasing NTP concentration or the duration of the transcription reaction. Finally, it is critical that template DNA barcoding is quantitative for both fractions of the transcription reaction. This is accomplished by the dUX-tagging procedure^31^, which is structured as a one-pot reaction to minimize sample handling variability and appends the fraction barcode to the constant P_RA1_ promoter sequence to eliminate the possibility of variant-dependent variability in barcoding efficiency. Together, the procedures described above enable the activity of RNA variants to be accurately measured using the cotranscriptional RNA-dependent DNA fractionation strategy.

Characterization of the *C. beijerinckii pfl* riboswitch consensus variant library revealed several pathways by which the ZTP aptamer can misfold and identified how stabilization of P1 base pairs can improve ZMP-responsiveness. The identification of these characteristics of *pfl* riboswitch folding required an approach in which classes of variants were isolated hierarchically. Base pair variation within the ZTP aptamer pseudoknot and terminator hairpin caused the most substantial effect on ZTP riboswitch function (Figure 3b, Supplementary Figure 2). This is unsurprising because ZTP riboswitches control transcription termination using a mechanism that requires helical switching between pseudoknot and terminator base pairs^25,40^. Once the effect of pseudoknot and terminator hairpin variability was isolated, it became clear that that the composition of nucleotides 46 and 43 in P1 contribute to ZMP-responsiveness (Figure 3f). However, rigorous evaluation of how nucleotides 46 and 43 contribute to ZMP-mediated transcription antitermination was only possible once a sub-population of variants that are prone to misfolding due to the presence of competitor helices was isolated (Figures 4 and 5). It is notable that the effect of nucleotide 46 and 43 identity and the presence of competitor helices was pervasive in variants with complete Watson-Crick pairing in both the pseudoknot and terminator hairpin. This indicates that variants within each pseudoknot and terminator composition class encounter the same cotranscriptional folding problems in addition to pseudoknot- and terminator-specific folding problems.

In the context of the *C*. *beijerinckii pfl* riboswitch scaffold, G24 variants exhibited a broad range of terminator efficiencies due to the apparent formation of an alternative pseudoknot (Fig. 3c). In the P1 subdomain, nucleotides that participate in the alternative pseudoknot partially overlap with nucleotides that participate in the native pseudoknot (Fig. 3a, d). In the P3 subdomain, nucleotides that participate in the alternative pseudoknot are adjacent to nucleotides that participate in the native pseudoknot, but emerge from RNAP first (Fig. 3a, d). In combination, the proximity of alternative pseudoknot base pairs to native pseudoknot base pairs and the earlier emergence of downstream alternative pseudoknot nucleotides from RNAP may cause G24 variants to be prone to aptamer misfolding. It is notable that the severity of G24-associated termination defects is affected by the identity of the 89:117 terminator base pair. This demonstrates that minor differences in expression platform sequence can have substantial context-dependent effects on riboswitch function.

The avoidance of competitor helices is an established mechanism for promoting accurate cotranscriptional RNA structure formation^60^ and has been shown to contribute to folding of the ZTP aptamer P3 subdomain^40^. Within the *C. beijerinckii pfl* riboswitch consensus variant library, we identified a class of variants that are prone to competitor helix-directed misfolding that inhibits both termination and ZMP-mediated antitermination (Figs. 4 and 5). In this case, ZTP aptamer misfolding required two distinct competitor helices that are mutually exclusive with P1 but not the native pseudoknot (Fig. 4b). Given that disrupting the native pseudoknot structure eliminated competitor helix-dependent terminator defects in most cases and that terminal PK and CH1 base pairs are adjacent, it is likely that CH1 inhibits terminator base pair propagation by directly stabilizing PK. It is unlikely that avoidance of the competitor helices identified here is a general property of ZTP riboswitch folding because CH1 comprises base pairs between conserved and non-conserved regions of the ZTP aptamer. However, these findings illustrate how competitor helix-directed misfolding can depend on the presence of several discrete structural elements.

The RNA-dependent DNA fractionation strategy used by TECdisplay experiments is conceptually similar to the mRNA display^61^ and ribosome display^62^ strategies for peptide evolution. In mRNA and ribosome display experiments, a nascent polypeptide is tethered to its mRNA template by covalent fusion or the ribosome complex, respectively. In this way, mRNA can be enriched based on the function of its encoded polypeptide. In TECdisplay experiments, template DNA is fractionated by the activity of its encoded RNA, which is physically tethered to its DNA template by the TEC. RNA-dependent DNA fractionation was previously used for *in vitro* selection by the DNA display method in which a run-off RNA transcript is tethered to the DNA molecule from which it was transcribed by a capture oligonucleotide that is covalently linked to the template DNA^63^. In principle, the procedures we have developed for TECdisplay could be used to perform cotranscriptional *in vitro* selection. For example, the strategy for separating DNA molecules on which a transcription termination event occurred from DNA molecules on which a terminator readthrough event occurred could potentially be applied to isolate riboswitches that regulate transcription. This approach would likely be synergistic with recent advances in methods for isolating synthetic riboswitches in which natural riboswitch aptamers are used as scaffold for *in vitro* evolution^22,64^.

## Online Methods

### Oligonucleotides

All oligonucleotides were purchased from Integrated DNA Technologies. A detailed description of all oligonucleotides including sequence, modifications, and purifications is presented in Supplementary Table 1. The sequence of the Ultramer oligonucleotide used in this study is presented in Supplementary Table 2.

### Proteins

Q5 DNA Polymerase, Q5U DNA Polymerase, Vent (exo-) DNA polymerase, *Sulfolobus* DNA Polymerase IV, *E*. *coli* RNA Polymerase Holoenzyme, RNase I_f_, Thermolabile Proteinase K, Thermolabile USER II Enzyme, T4 DNA Ligase, ET SSB, and Lambda Exonuclease were purchased from New England Biolabs (NEB). UltraPure BSA was purchased from Invitrogen. GreB was a gift from R. Landick (University of Wisconsin, Madison).

### DNA template preparation

Supplementary Table 3 provides details for the oligonucleotides and processing steps used for every DNA template preparation used in this work. Supplementary Table 4 provides DNA template sequences.

Because the maximum Ultramer length is 200 nt, it was necessary to amplify DNA templates for TECdisplay in two steps. The first PCR, which appended sequence from the P_RA1_ promoter, was performed as a 100 μl reaction containing 1X Q5 Reaction Buffer (NEB), 1X Q5 High GC Enhancer (NEB), 200 μM dNTPs (NEB), 250 nM TECD020.PRA1_C3SC1.F (Supplementary Table 1), 250 nM TECD017.dRP1_NoMod.R (Supplementary Table 1), 0.02 nM ULT004.C3SC1_pflCONSv1_VRA3 template (Supplementary Table 2), and 0.02 U/μl Q5 DNA polymerase (NEB). Amplification was performed using the program: 95 °C for 3 min, [95 °C for 20s, 65 °C for 30s, 72 °C for 20s] x 12 to 15 cycles, 72 °C for 5 min. PCR products were mixed with an equal volume of SPRI beads, purified as described below in the section *Purification of DNA by solid-phase reversible immobilization (SPRI)*, and eluted into 20 μl of 10 mM Tris-HCl (pH 8.0). DNA was quantified using the Qubit dsDNA HS Assay Kit (Invitrogen) with a Qubit 4 Fluorometer (Invitrogen). Molarity was determined using the length of the DNA.

DNA templates that contained an internal etheno-dA transcription stall site were PCR amplified from the purified dsDNA prepared above. This second PCR was performed as four 100 μl reactions containing 1X Q5U Reaction Buffer (NEB), 1X Q5 High GC Enhancer, 200 μM dNTPs, 500 nM TECD007.pRA1_2dU.F (Supplementary Table 1), 500 nM TECD002.dRP1iEthDA.R (Supplementary Table 1), 1 μl of 1 pM LPRA5_C3SC1_ULT004_dRP1 template DNA (Supplementary Table 4), and 0.02 U/μl Q5U DNA polymerase (NEB). Amplification was performed as a 2-step PCR using the program: 98 °C for 30s, [98 °C for 10s, 72 °C for 2 min] x 30 cycles, 72 °C for 1 min. After the reaction was complete, the thermal cycler was held at 72C, 33 μl of the original PCR was diluted into 66 μl of fresh PCR master mix that contained 1X Q5U Reaction Buffer, 1X Q5 High GC Enhancer, 200 μM dNTPs, 750 nM TECD007.pRA1_2dU.F (Supplementary Table 1), 750 nM TECD002.dRP1iEthDA.R (Supplementary Table 1), and 0.02 U/ μl Q5U DNA polymerase. These diluted reactions were then amplified for two additional cycles.

PCR products were ethanol precipitated by mixing with 3 volumes of 100% ethanol, 0.1 volumes of 3 M sodium acetate (pH 5.5), and 1.5 μl of GlycoBlue Coprecipitant (Invitrogen) and chilling at −80°C for at least 30 minutes. The samples were centrifuged at 18,500 x g and 4°C for 30 minutes, the supernatant was removed, the samples were centrifuged again briefly to pull down residual ethanol, and residual ethanol was removed. The pellets were washed by adding 500 μl of 70% ethanol and centrifuged at 18,500 x g and 4 °C for 2 minutes. The supernatant was removed, the samples were centrifuged again briefly to pull down residual ethanol, and residual ethanol was removed. The pellets were dissolved in 100 μl of 10mM Tris-HCl (pH 8.0) and mixed with 20 μl of 6X DNA Loading Dye [10 mM Tris-HCl (pH 8.0), 30% (v/v) glycerol, 0.48% (w/v) SDS, 0.05% (w/v) Bromophenol Blue]. The sample was run on a 8% native TBE-polyacrylamide gel at 120V for 75 minutes. Gels were stained with SYBR Gold Nucleic Acid Stain (Invitrogen) for 10 minutes. The modified DNA template of interest was visualized using a blue light transilluminator and cut out of the gel using a clean razor blade. The gel slice was placed inside a 600 μl microcentrifuge tube with a needle hole in the bottom, which was placed in a 2 ml collection tube and centrifuged at 18,500 x g and 4°C for 5 minutes to crush the gel. The gel was resuspended in 3 μl of Polyacrylamide Gel Extraction Buffer (300 mM sodium acetate [pH 5.5], 1 mM EDTA) per mg of gel and incubated at room temperature with rotation (∼15 rpm) overnight. The sample was centrifuged at 18,500 x g and 4°C for 5 minutes and the supernatant was transferred to a Spin-X 0.22 μm cellulose acetate centrifuge tube filter. The gel was resuspended in 1 μl of Polyacrylamide Gel Extraction Buffer per mg of gel, mixed by vortexing, centrifuged at 18,500 x g and 4°C for 5 minutes, and the supernatant was transferred to the filter column. Samples were centrifuged at 2000 x g and 4°C for 3 minutes and the filtrate was collected. The filtrate was mixed with an equal volume of phenol:chloroform:isoamyl alcohol (25:24:1, v/v), mixed by vortexing and inversion, and centrifuged at 18,500 x g and 4°C for 5 minutes. The aqueous phase was collected and transferred to a new tube. The DNA was ethanol precipitated as described above and resuspended in 100 μl of 10 mM Tris-HCl (pH 8.0).

The purified DNA template was then completed using translesion primer extension, which extends the non-transcribed DNA strand beyond the etheno-dA modification in the template DNA strand^29^. Translesion DNA synthesis using *Sulfolobus* DNA polymerase IV (NEB) was performed as two 100 μl reactions exactly as described previously^29,65^. The resulting dsDNA was mixed with an equal volume of SPRI beads and purified as described below in the section *Purification of DNA by solid-phase reversible immobilization (SPRI)*. Following purification, the resulting DNA template was quantified using the Qubit dsDNA HS Assay Kit (Invitrogen) with a Qubit 4 Fluorometer and molarity was determined using the length of the DNA.

### Purification of DNA by solid-phase reversible immobilization (SPRI)

SPRI beads were prepared in-house using the ‘DNA Buffer’ variation of the procedure by Jolivet and Foley^66^. Samples were mixed with a variable amount of SPRI beads (depending on the procedure, details are in each relevant section), incubated at room temperature for 5 min, and placed on a magnetic stand for 3 min so that the beads collected on the tube wall. The supernatant was aspirated and discarded, and the beads were washed twice by adding a volume of 70% ethanol at least 200 μl greater than the combined volume of the sample and SPRI beads to the tube without disturbing the bead pellet while it remained on the magnetic stand. The samples were incubated at room temperature for 1 min before aspirating and discarding the supernatant. Residual ethanol was evaporated by placing the open microcentrifuge tube in a 37°C dry bath for ∼15 s with care taken to ensure that the beads did not dry out. Purified DNA was eluted by resuspending the beads in a variable amount of 10 mM Tris-HCl (pH 8.0) (depending on the procedure, details are in each relevant section), allowing the samples to sit undisturbed for 3 min, placing the sample on a magnetic stand for 1 min so that the beads collected on the tube wall, and transferring the supernatant, which contained purified DNA, into a screw-cap tube with an O-ring.

### TECdisplay

#### Preparation of streptavidin-coated magnetic beads

5 μl of 10 mg/ml Dynabeads MyOne Streptavidin C1 beads (Invitrogen) per 25 μl sample volume were prepared in bulk exactly as described previously^28,52^. The resulting beads were resuspended at a concentration of 2 μg/μl in Buffer TX (1X transcription buffer [20 mM Tris-HCl (pH 8.0), 50 mM KCl, 1 mM DTT, 0.1 mM EDTA (pH 8.0)] supplemented with 0.1% Triton X-100), split into in 25 μl aliquots, and stored on ice until use.

#### Preparation of ^C3-SC1^TECs for quantitative in vitro transcription

Quantitative *in vitro* transcription was performed by isolating magnetic bead-immobilized, synchronized TECs using the C3-SC1 leader (^C3-SC1^TECs) essentially as described previously^28^. A detailed protocol for the preparation of ^C3-SC1^TECs is available^52^. The original method for ^C3-SC1^TECs purification with application-specific modifications is presented below. One sample volume is defined as 25 μl.

*In vitro* transcription reactions containing 1X Transcription Buffer, 0.1 mg/ml BSA, 100 μM ApU dinucleotide, 1X UGC Start NTPs (25 μM UTP, 25 μM GTP, 25 μM CTP [Cytiva, cat. no. 27202501]), 20 nM PRA1_2dU_C3SC1_ULT004_dRP1_iEthdA template DNA, 0.032 U/μl *E*. *coli* RNAP holoenzyme (NEB), and 25 nM TECD018.Cap3_3BioTEG oligonucleotide (Supplementary Table 1) were prepared in bulk on ice; at this point the total reaction volume per sample was 20 μl due to the omission of 10X (200 μg/ml) heparin (Millipore Sigma, catalog no. H5515) and 10X Start Solution (100 mM MgCl_2_, 100 μg/ml rifampicin). The bulk transcription reaction was placed in a dry bath set to 37°C for 20 minutes to form open promoter complexes. After 20 minutes, 2.5 μl of 200 μg/ml heparin per sample volume was added to the reaction, and the sample was mixed by pipetting and incubated at 37°C for 5 minutes to sequester free RNAP and enrich for heparin-resistant open promoter complexes; the final concentration of heparin was 20 μg/ml. After 5 minutes, 2.5 μl of room temperature 10X Start Solution per sample volume was added to the transcription reaction for a final concentration of 10 mM MgCl_2_ and 10 μg/ml rifampicin. The transcription reaction was mixed by pipetting and incubated at 37°C for 20 minutes to walk RNAP to the C_+31_ synchronization site in the C3-SC1 leader and hybridize the TECD018.Cap3_3BioTEG oligonucleotide (Supplementary Table 1) to nascent RNA. At this time, 25 μl of ∼2 μg/μl pre-equilibrated streptavidin-coated magnetic beads per sample volume, Wash Buffer (1X Transcription Buffer, 10 mM MgCl_2_, and 0.05% Tween-20), Elution Buffer 1 (1.28X Transcription Buffer, 0.1mg/ml BSA, 12.82 mM MgCl_2_, 10 μg/ml Rifampicin, and 100nM GreB [a gift from R. Landick, UW Madison]), and Elution Buffer 2 (1X Transcription Buffer, 0.1mg/ml BSA, 10 mM MgCl_2_, and 10 μg/ml Rifampicin) were placed at room temperature.

After ∼18 minutes, the magnetic beads were placed on a magnetic stand and storage buffer was removed. After 20 minutes, the pre-equilibrated streptavidin-coated magnetic beads were resuspended with the bulk transcription reaction by pipetting and incubated at room temperature with rotation (∼15 rpm) for 15 minutes to immobilize ^C3-SC1^TECs. After 15 minutes, the bead binding mixture was spun briefly in a Labnet Prism mini centrifuge by quickly flicking the switch on and off so that liquid was removed from the tube cap, but the speed of the mini centrifuge remained as low as possible. The sample was placed on a magnet stand for 1 minute and the supernatant was discarded. The 1.7 ml tube containing the sample was removed from the magnet stand and the beads were gently resuspended in 1 ml of room temperature Wash Buffer and incubated at room temperature with rotation for 5 minutes. The sample was placed on a magnet stand for 1 minute and the supernatant was discarded. Immobilized ^C3-SC1^TECs were gently resuspended in 22 μl of room temperature Elution Buffer 1 per sample volume.

#### Termination-dependent partitioning of template DNA

The ^C3-SC1^TECs and 1.7 ml microcentrifuge tubes containing ZMP Chase Mix were placed in a dry bath set to 37°C for 2 minutes to prewarm. For samples that contained ≤1 mM ZMP, ZMP Chase Mix contained 2.5 μl of 5 mM NTPs (Cytiva, cat. no. 27202501) and 0.5 μl of 50X ZMP solution in DMSO or, in the case of 0 mM ZMP samples, 0.5 μl of DMSO. For the 3.16 mM ZMP sample, ZMP Chase Mix contained 2.5 μl of 5 mM NTPs and 1.58 μl of 50 mM ZMP. The final NTP concentration for all samples was 500 μM. Because the 50 mM ZMP stock was dissolved in DMSO, samples that contained ≤1 mM ZMP also contained 2% DMSO; the 3.16 mM sample contained 6.32% DMSO. After 2 minutes, 22 μl (≤1 mM ZMP samples) or 20.9 μl (3.16 mM sample) of immobilized ^C3-SC1^TECs were mixed with ZMP Chase Mix and incubated at 37°C for 3 minutes. The transcription elongation factor GreB was included in this reaction (as part of Elution Buffer 1) to rescue backtracked elongation complexes, which could potentially cause experimental noise because DNA that contains a backtracked TEC will always partition into the pellet fraction. After 3 minutes of transcription, samples were placed on a magnetic stand for 1 minute and the supernatant was transferred to a 200 μl thin-walled PCR tube; this is the unbound (UNB) fraction. The remaining beads were gently resuspended in 25 μl of Elution Buffer 2 and transferred to a 200 μl thin-walled PCR tube; this is the bound (BND) fraction. Samples remained on ice until RNA degradation.

#### RNA degradation

0.5 μl of RNase I_f_ (NEB) was added to each 25 μl transcription reaction fraction and mixed by pipetting. Samples were incubated on a thermal cycler set to 37°C for 15 minutes. Samples were then incubated at 70°C for 20 minutes to heat-inactivate RNase I_f_. The bound fraction was placed on a magnetic stand for 1 minute, and the supernatant, which contains eluted template DNA, was transferred to a new 200 μl thin-walled PCR tube.

#### Protein degradation

1 μl of Thermolabile Proteinase K (NEB) was added to each 25.5 μl sample and mixed by pipetting. Samples were incubated on a thermal cycler set to 37°C for 30 minutes. Samples were then incubated at 65°C for 20 minutes to heat-inactivate Thermolabile Proteinase K. At this point samples were either barcoded as described below or stored at −20°C overnight before barcoding.

#### Fraction barcoding by deoxyuridine excision tagging (dUX-tagging)

Template DNA was barcoded using a modified version of the dUX-tagging procedure, which quantitatively appends a barcode and Illumina adapter to dsDNA in a one-pot reaction. USER digestions were prepared on ice by adding 3 μl of 10X T4 DNA Ligase Buffer (NEB) to each 26.5 μl sample, pipetting to mix, then adding 0.5 μl of Thermolabile USER II Enzyme (NEB) and pipetting to mix again. Samples were incubated at 37°C for 30 minutes on a pre-warmed thermal cycler block with a heated lid set to 45°C and held at 12°C after the incubation was complete. Following USER digestion, 0.4 μl of 500 ng/μl (200 ng) ET SSB (NEB) and 0.6 μl of 2.5 μM (1.5 pmol) of the TECD021.5pBND_pRA1m12_VRA5 or TECD022.5pUNB_pRA1m12_VRA5 oligonucleotides (for bound and unbound fractions, respectively; Supplementary Table 1) were added to each sample and mixed by pipetting. To anneal the tagging oligo and inactivate Thermolabile USER II Enzyme, the master mix was placed on a thermal cycler block set to 70°C with a heated lid set to 105°C and slowly cooled using the protocol: 70°C for 5 minutes, ramp to 65°C at 0.1°C/s, 65°C for 5 minutes, ramp to 60°C at 0.1°C/s, 60°C for 2 minutes, ramp to 37°C at 0.1°C/s, hold at 37°C. After annealing the tagging oligo, 1 μl of T4 DNA ligase (NEB) was added to each sample and the samples were mixed by pipetting while the reaction remained at 37°C. The ligation reactions were incubated at 37°C for 5 minutes and then at 65°C for 10 minutes to heat-inactivate T4 DNA ligase. A primer extension reaction containing the 32 μl tagged DNA sample (which at this point contains a 5’ overhang), 1X ThermoPol Buffer (NEB), 0.2 mM dNTPs, 2.5% formamide, and 0.02 U/μl Vent (exo-) DNA Polymerase (NEB) was prepared on ice for each sample. These 100 μl primer extension reactions were placed on a pre-heated 72°C thermal cycler block for 5 minutes and then moved to an ice-cold aluminum PCR tube block to snap cool. At this point the template DNA is fully barcoded, and the bound and unbound fractions of each sample were combined. The barcoded DNA was then mixed with 200 μl of SPRI beads and purified as described in above in the section *Purification of DNA by solid-phase reversible immobilization (SPRI)* to deplete excess tagging oligonucleotide. Purified DNA was eluted into 50 μl of 10 mM Tris-HCl (pH 8.0).

#### Non-transcribed DNA strand degradation

Following template DNA barcoding and purification, the non-transcribed DNA strand was selectively degraded using lambda exonuclease so that sequencing libraries are amplified exclusively from the transcribed DNA strand, which is guaranteed to correspond to the transcribed RNA sequence. This ensures that TECdisplay measurements are not confounded by heteroduplexes, which may be present in the DNA template preparation. Lambda exonuclease reactions were prepared on ice in 200 μl thin-walled PCR tubes by adding 5.56 μl of 10X Lambda Exonuclease Reaction Buffer (NEB) to the 50 μl purified DNA samples, pipetting to mix, adding 0.5 μl of Lambda Exonuclease (NEB), and pipetting to mix again. Samples were incubated at 37°C on a thermal cycler block for 5 minutes, and then at 75°C for 10 minutes to heat-inactivate lambda exonuclease.

#### Purification of ssDNA by phenol-chloroform extraction

Following non-transcribed DNA strand degradation, the volume of each sample was raised to 150 μl by adding 94 μl of 10mM Tris-HCl (pH 8.0). Samples were mixed with an equal volume (150 μl) of Tris (pH 8) buffered phenol:chloroform:isoamyl alcohol (25:24:1, v/v), mixed by vortexing and inversion, and centrifuged at 18,500 x g and 4°C for 5 minutes. The aqueous phase was collected and transferred to a new tube. ssDNA was precipitated by mixing the aqueous phase with 450 μl of 100% ethanol, 15 μl of 3M sodium acetate (pH 5.5), and 1.5 μl of GlycoBlue Coprecipitant and chilling at −80°C for at least 30 minutes. The samples were centrifuged at 18,500 x g and 4°C for 30 minutes, the supernatant was removed, the samples were centrifuged again briefly to pull down residual ethanol, and residual ethanol was removed. The pellets were washed by adding 500 μl of 70% ethanol and centrifuged at 18,500 x g and 4 °C for 2 minutes. The supernatant was removed, the samples were centrifuged again briefly to pull down residual ethanol, and residual ethanol was removed. The pellets were dissolved in 100 μl of 10mM Tris-HCl (pH 8.0).

### Test amplification of TECdisplay libraries

The number of PCR amplification cycles needed for TECdisplay libraries was determined by performing a test amplification adapted from Mahat et al.^67^. 20 μl PCRs contained 14 μl of Q5 PCR master mix and 6 μl of a 16-fold or 64-fold dilution of the ssDNA libraries, such that the final concentration of the reaction components were 1X Q5 Reaction Buffer, 1X Q5 High GC Enhancer, 200 μM dNTPs, 250 nM RPIX (Supplementary Table 1), 250 nM TECD017.dRP1_NoMod.R (Supplementary Table 1), and 0.02 U/μl Q5 DNA Polymerase. One PCR for each ssDNA dilution was amplified for 11 or 12 cycles using the program: 98 °C for 30s, [98 °C for 10s, 62 °C for 20s, 72 °C for 20s] x 11 or 12 cycles, 72 °C for 2 min. This format yields test amplifications that correspond to 4, 5, 6, and 7 cycles in the final library amplification described below. Following amplification, each PCR was mixed with 4 μl of 6X DNA Loading Dye and run on a native TBE-polyacrylamide gel to determine the number of cycles needed to amplify the libraries for high-throughput sequencing.

### Preparation of DNA libraries for sequencing

Indexed dsDNA libraries were prepared for Illumina sequencing by amplifying each sample in a 50 μl PCR that contained 1X Q5 Reaction Buffer, 1X Q5 High GC Enhancer, 200 μM dNTPs, 500 nM RPI Indexing Primer (Supplementary Table 1), 500 nM TECD017.dRP1_NoMod.R (Supplementary Table 1), 12 μl of ssDNA library, and 0.02 U/μl Q5 DNA Polymerase. Amplification was performed using the program 98 °C for 30s, [98 °C for 10s, 62 °C for 20s, 72 °C for 20s] x [# of cycles], 72 °C for 2 min, where “# of cycles” corresponds the optimal number of cycles determined by test amplification. Following amplification, 50 μl PCRs were mixed with 100 μl of SPRI beads and purified as described in *Purification of DNA by solid-phase reversible immobilization (SPRI)*. DNA was eluted into 20 μl of 10 mM Tris-HCl (pH 8.0), mixed with 40 μl of SPRI beads, and purified a second time. Twice-purified DNA was eluted into 10 μl of 10 mM Tris-HCl (pH 8.0) and quantified using the Qubit dsDNA HS Assay Kit (Invitrogen) with a Qubit 4 Fluorometer.

### High-throughput DNA sequencing

Sequencing of TECdisplay libraries was performed by Novogene Co. on an Illumina HiSeq 4000 System using 2x150 PE reads with 35% PhiX spike in. TECdisplay libraries were sequenced at a depth of ∼200 to ∼250 million PE reads. Due to the extremely low complexity of TECdisplay libraries, the best sequencing outcomes were obtained by pooling TECdisplay libraries with libraries of a different type (e.g. TECprobe-VL^38^ libraries).

### TECdisplay data analysis

Source code and documentation for custom data analysis tools is available at https://github.com/e-strobel-lab/TECtools. Target sequences used to map sequencing reads were generated by `variant_maker`. Sequencing read pre-processing and alignment was performed by `TECdisplay_mapper`, which manages sequencing read preprocessing using `fastp`^68^ and maps reads to targets that were generated by `variant_maker`. During preprocessing, `fastp` trims adapter sequences (which are rarely present in the sequencing reads), performs error correction in overlapping regions of read pairs, merges read pairs, and extracts the fraction barcode from the head of read 2. `TECdisplay_mapper` then maps merged sequencing reads to targets by identifying the location of the SC1 hairpin in the sequencing read, generating a key from bases at locations within the read that are expected to contain variable bases, using the key to search a hash table of all target sequences for a potential match, and verifying the match by comparing the target RNA sequence from the sequencing read to the potential target match. If a match to a reference target is identified, the read is assigned to the bound or unbound fraction based on its fraction barcode. After read mapping is complete, the fraction of bound reads for each variant is calculated by dividing the number of bound reads by the sum of bound and unbound reads. During interpretation of the data, variants within each data set were filtered by nucleotide and base pair identity using `TECdisplay_navigator` or `TECdisplay_Hnav`, which coordinates hierarchical `TECdisplay_navigator` analyses. Detailed instructions for using all of the software described above are available at https://github.com/e-strobel-lab/TECtools.

### Characterization of RNA-dependent DNA partitioning accuracy

To assess the accuracy of RNA-dependent DNA partitioning, ^C3-SC1^TECs were prepared as described above in the section *Preparation of* ^C3-SC1^TECs *for quantitative in vitro transcription* using the PRA1_2dU_C3SC1_CbePfl_dRP1_iEthdA DNA template (Supplementary Table 4), which encodes the wild-type *Cbe pfl* ZTP riboswitch. After ^C3-SC1^TEC purification, transcription was resumed by adding NTPs to a final concentration of either 100 μM or 500 μM in the presence and absence of 1 mM ZMP and template DNA was partitioned as described above in the section *Termination-dependent partitioning of template DNA*. Upon collection, the 25 μl supernatant of each sample was mixed with 125 μl of Stop Solution (0.6 M Tris-HCl [pH 8.0], 12 mM EDTA). Each bead pellet was resuspended in 25 μl of Formamide Elution Solution (95% formamide (v/v), 10 mM EDTA) and boiled at 100 °C for 5 minutes. The pellet samples were placed on a magnetic stand and each 25 μl supernatant was mixed with 125 μl of Stop Solution. The 150 μl samples were mixed with an equal volume of Tris (pH 8) buffered phenol:chloroform:isoamyl alcohol (25:24:1, v/v) and purified by phenol/chloroform extraction and ethanol precipitation as described above in the section *Purification of ssDNA by phenol-chloroform extraction* except that the resulting pellets were resuspended in 15 μl of formamide loading dye (90% (v/v) deionized formamide, 1X transcription buffer, 0.01% (w/v) bromophenol blue) and assessed by denaturing PAGE as described in the section *Denaturing Urea-PAGE*. Quantification of band intensity was performed using ImageJ 1.53k exactly as described previously^44^. For DNA bands, fraction terminator readthrough was calculated by dividing the intensity of the pellet band by the sum of the intensities of the supernatant and pellet bands. For RNA bands, band intensity was normalized by RNA length and fraction terminator readthrough was calculated by dividing the normalized intensity of the readthrough band by the sum of the normalized intensities of the terminated and readthrough bands, as described previously^40^.

### Intermediate fraction analysis of the dUX-tagging procedure

To visualize intermediate products of the optimized dUX-tagging procedure, the PRA1_2dU_C3SC1_CbePfl_dRP1 DNA template (Supplementary Table 4) was barcoded in bulk using the procedure described above in the section *Fraction barcoding by deoxyuridine excision tagging (dUX-tagging)* and intermediate fractions were collected after key processing steps. Note that the volume of each fraction changes to account for the increase in sample volume as reagents are added throughout the dUX-tagging procedure. The input DNA fraction was collected by transferring 24.5 μl of the reaction to 125 μl of stop solution before USER enzyme was added. The USER-digested fraction was collected by transferring 25 μl of the reaction to 125 μl of stop solution after USER digestion. For experiments in which the tagging oligo ligation was performed at 37 °C for 5 min, which contained ET SSB during the ligation, the post-ligation fraction was collected by transferring 26.9 μl of the reaction to 125 μl of Stop Solution. For experiments in which the tagging oligo ligation was performed at 25 °C for 60 min, which did not contain ET SSB during the ligation, the post-ligation fraction was collected by transferring 26.5 μl of the reaction to 125 μl of Stop Solution. The post-primer extension fraction was collected by mixing the 100 μl primer extension reaction with 50 μl of stop solution supplemented with 1.8 μl of 500 mM EDTA (pH 8.0) so that the final EDTA concentration was 10 mM. Primer extension reactions were mixed with 200 μl of SPRI beads and purified as described in above in the section *Purification of DNA by solid-phase reversible immobilization (SPRI)*. Purified DNA was eluted into 50 μl of 10 mM Tris-HCl (pH 8.0) and one sample was collected as the post-purification fraction by mixing with 100 μl of Stop Solution. The final sample was treated with lambda exonuclease as described in the section *Non-transcribed DNA strand degradation*, and the ∼56 μl ssDNA product fraction was mixed with 94 μl 10 mM Tris-HCl (pH 8). Fractions were mixed with an equal volume (150 μl) of Tris (pH 8) buffered phenol:chloroform:isoamyl alcohol (25:24:1, v/v) and processed by phenol/chloroform extraction and ethanol precipitation as described above in the section *Purification of ssDNA by phenol-chloroform extraction.* Following ethanol precipitation, each pellet was resuspended in 15 μl of formamide loading dye. The samples were analyzed by denaturing PAGE as described in the section *Denaturing Urea-PAGE*.

### Optimization of dUX-tagging ligation time

Eight 25 μl dUX-tagging reactions containing 1X T4 DNA Ligase Buffer, 5 nM PRA1_2dU_C3SC1_CbePfl_dRP1 template DNA (Supplementary Table 4), and 0.02 U/μl Thermolabile USER II Enzyme were prepared in bulk in thin-walled 200 μl PCR tubes on ice; One 24.5 μl reaction volume was transferred to 125 μl of Stop Solution for collection as an input DNA fraction before USER enzyme was added. USER digestion was performed as described in the section *Fraction barcoding by deoxyuridine excision tagging (dUX-tagging)* and one reaction volume (25 μl) was transferred to 125 μl of Stop Solution for collection as the USER-digested sample and kept on ice. 0.5 μl of 2.5 μM TECD021.5pBND_pRA1m12_VRA5 or TECD022.5pUNB_pRA1m12_VRA5 (Supplementary Table 1) per reaction volume and 0.4 μl of ET SSB per reaction volume were added to the master mix. The tagging oligo was annealed as described in the section *Fraction barcoding by deoxyuridine excision tagging (dUX-tagging)* and the sample was held at 37 °C. One 25.9 μl fraction was collected as a zero time point before T4 DNA ligase was added. The master mix was aliquoted into five separate thin-walled 200 μl PCR tubes. Each sample was mixed with 1 μl of T4 DNA ligase and incubated at 37 °C for the indicated amount of time (30 s, 1 min, 2 min, 5 min, or 10 min) before being incubated at 65°C for 10 minutes to heat-inactivate T4 DNA ligase. After heat inactivation, samples were mixed with 125 μl of Stop Solution and kept on ice. The samples were phenol/chloroform extracted and ethanol precipitated as described in the section *Purification of ssDNA by phenol-chloroform extraction* and each pellet was resuspended in 15 μl of formamide loading dye. The samples were analyzed by denaturing PAGE as described in the section *Denaturing Urea-PAGE*.

### Optimization of lambda exonuclease digestion time

Seven 25 μl dUX-tagging reactions were prepared in bulk as described in the section *Optimization of dUX-tagging ligation time* and processed as described in the section *Fraction barcoding by deoxyuridine excision tagging (dUX-tagging)* through SPRI bead purification; One 24.5 μl reaction volume was transferred to 125 μl of Stop Solution for collection as an input DNA fraction before USER enzyme was added to the initial master mix, and a second 50 μl sample was transferred to 100 μl of Stop Solution after SPRI bead purification for collection as the pre-lambda exonuclease digestion fraction. Each of the remaining five samples was mixed with 5.56 μl of Lambda Exonuclease Reaction Buffer and 0.5 μl of Lambda Exonuclease and incubated at 37°C for the indicated amount of time (30 s, 1 min, 2 min, 5 min, or 10 min) before being mixed with 1.1 μl of 0.5 M EDTA and incubated at 75°C for 10 minutes to heat inactivate lambda exonuclease. Samples were phenol/chloroform extracted and ethanol precipitated as described in the section *Purification of ssDNA by phenol-chloroform extraction* and each pellet was resuspended in 15 μl of formamide loading dye. The samples were analyzed by denaturing PAGE as described in the section *Denaturing Urea-PAGE*.

### Denaturing Urea-PAGE

Denaturing Urea-PAGE was performed exactly as described previously^44^. Briefly, samples in formamide loading dye were heated at 95°C for 5 min, and snap-cooled on ice for 2 min. Denaturing Urea-PAGE was performed using 8 or 10% gels prepared using the SequaGel UreaGel 19:1 Denaturing Gel System (National Diagnostics) for a Mini-PROTEAN Tetra Vertical Electrophoresis Cell as described previously^44^. Gels were stained with SYBR Gold Nucleic Acid Stain (Invitrogen) and scanned using a Typhoon RGB biomolecular imager or an Azure Sapphire biomolecular imager.

## Data availability

Raw sequencing data that support the findings of this study have been deposited in the Sequencing Read Archive (https://www.ncbi.nlm.nih.gov/sra) with the BioProject accession code PRJNA976257. Individual BioSample accession codes are available in Supplementary Table 5. Fully processed TECdisplay data have been deposited in Zenodo (<u>10.5281/zenodo.10412935</u>). Source gel images have been deposited in Zenodo (<u>10.5281/zenodo.10425136</u>)

## Code availability

TECtools can be accessed at https://github.com/e-strobel-lab/TECtools/releases/tag/v1.1.0.

## Supplementary information

This article contains supplementary information.

## Author Contributions

E.J.S., conceptualization; E.J.S., methodology; S.L.K., investigation; S.L.K. and E.J.S, Validation; S.L.K. and E.J.S, Formal Analysis; E.J.S, Software; S.L.K. and E.J.S, writing – original draft; S.L.K. and E.J.S, writing – review & editing; E.J.S., supervision; E.J.S., funding acquisition.

## Funding

This work was supported by the National Institute of General Medical Sciences of the National Institutes of Health under Award Number R35GM147137 (to E.J.S) and by start-up funding from the University at Buffalo (to E.J.S). The content is solely the responsibility of the authors and does not necessarily represent the official views of the National Institutes of Health.

## Conflict of Interest

The authors have no conflicts of interest with the contents of this article.

## Supplementary Figures

**Supplementary Figure 1.**
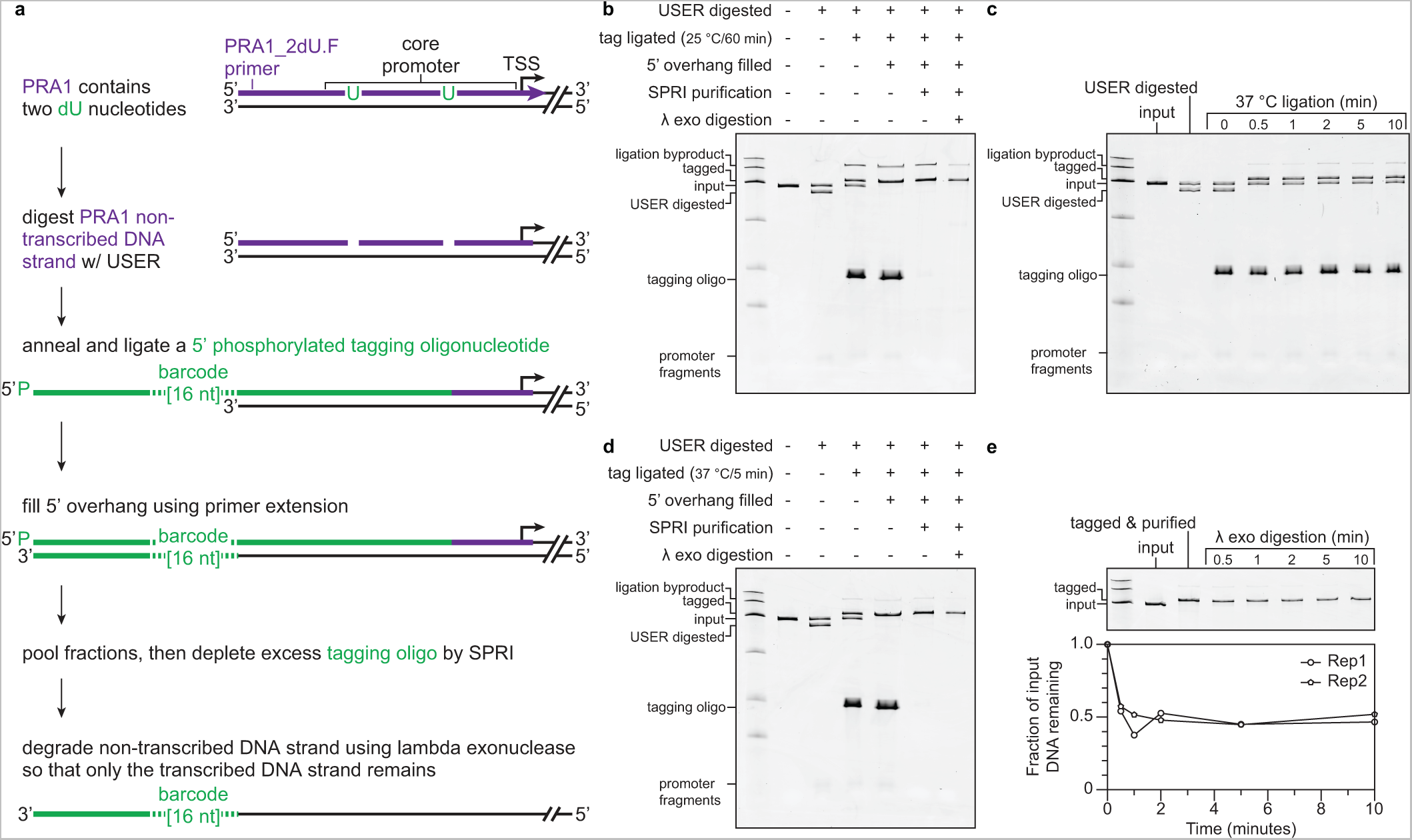
Optimization of dU-excision tagging reaction parameters. **a,** Overview of the dU-excision tagging procedure. Two dU nucleotides are excised from the P_RA1_ promoter. The 5’-phosphorylated tagging oligonucleotide is then annealed in place of the fragmented promoter strand and ligated to the template DNA. The resulting 5’ overhang is then filled in by primer extension. Tagged DNA from separate TECdisplay fractions is then pooled together and excess tagging oligonucleotide is depleted using SPRI beads. The non-transcribed DNA strand, which contains a 5’ phosphate from the tagging oligonucleotide, is then selectively degraded using lambda exonuclease so that only the transcribed DNA strand remains. **b**, Visualization of dUX-tagging reaction intermediates using a 5’-phosphorylated tagging oligonucleotide and the ligation conditions used in the original protocol (25 °C for 60 minutes). In these conditions, the 5’ phosphorylated tagging oligo causes a ligation byproduct to form. **c**, Optimization of the dUX-tagging ligation. A dUX-tagging ligation time course was performed at 37 °C. 5 minutes was selected as the reaction duration used in the optimized procedure. **d**, Visualization of dUX-tagging reaction intermediates using a 5’-phosphorylated tagging oligonucleotide and the optimized ligation conditions (37 °C for 5 minutes). In these conditions, formation of the ligation byproduct observed in panel b is reduced. **e**, Optimization of lambda exonuclease digestion reaction time. The lambda exonuclease digestion is complete within 5 minutes and the resulting ssDNA, which does not contain a 5’ phosphate, is stable. 5 minutes was selected as the reaction duration used in the optimized procedure. All gels are n=2. dU, deoxyuridine; USER, Uracil-Specific Excision Reagent; SPRI, solid-phase reversible immobilization.

**Supplementary Figure 2.**
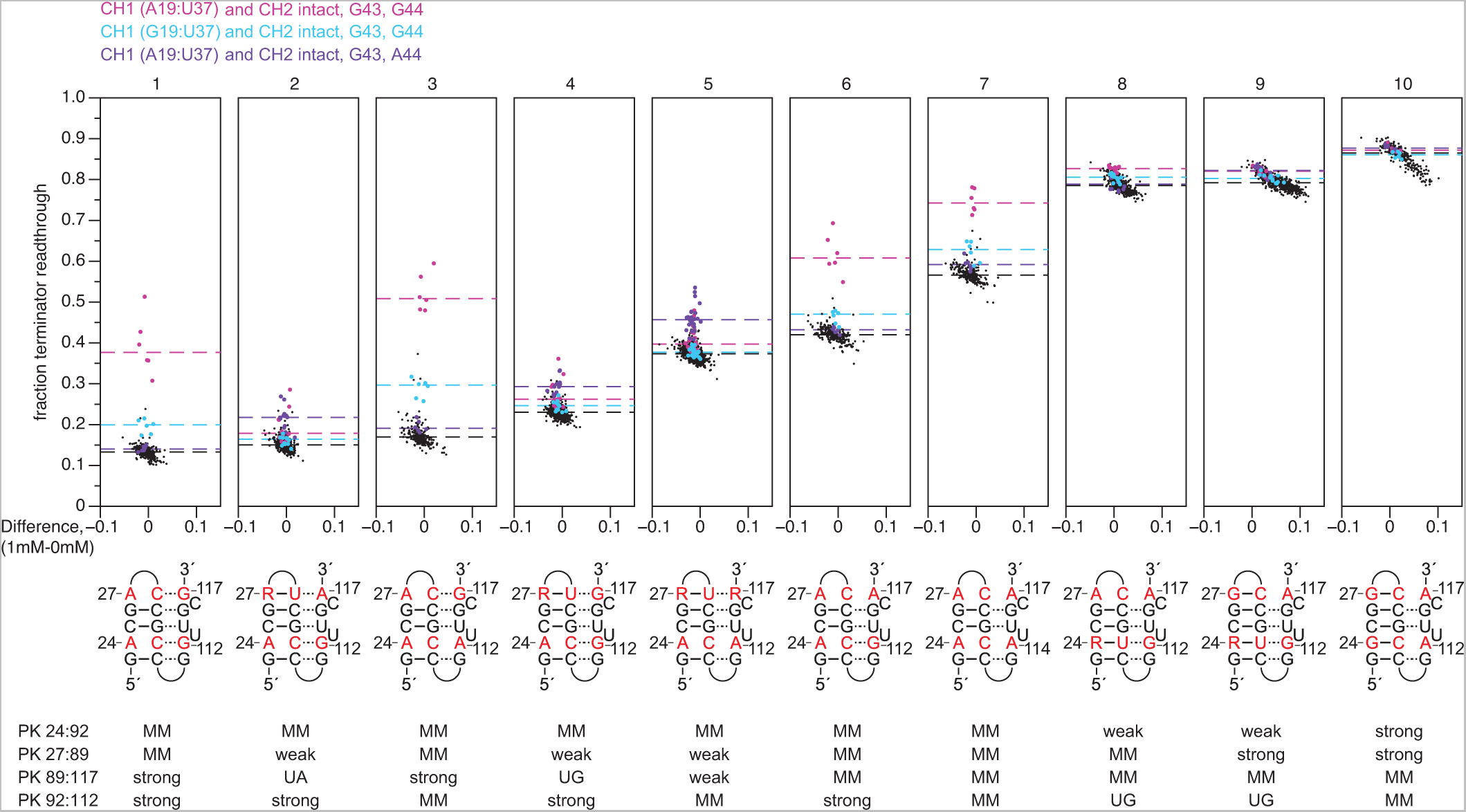
Perturbation of *C. beijerinckii pfl* ZTP riboswitch pseudoknot and terminator base pairs. Plots showing the effect of pseudoknot and terminator perturbations on fraction terminator readthrough. Variants are distributed across the x-axis by the difference in fraction terminator readthrough measured for the 1 mM ZMP and 0 mM ZMP conditions. Variants that can form the CH1 and CH2 competitor helices and have the capacity to form an alternative pseudoknot as depicted in Figure 4f are highlighted in magenta, cyan, or purple. The median fraction terminator readthrough for each group of variants is shown as a horizontal dashed line. All data are from the experiment shown in Figure 2. CH1, competitor helix 1; CH2, competitor helix 2; PK, pseudoknot; MM, mismatch.

**Supplementary Figure 3.**
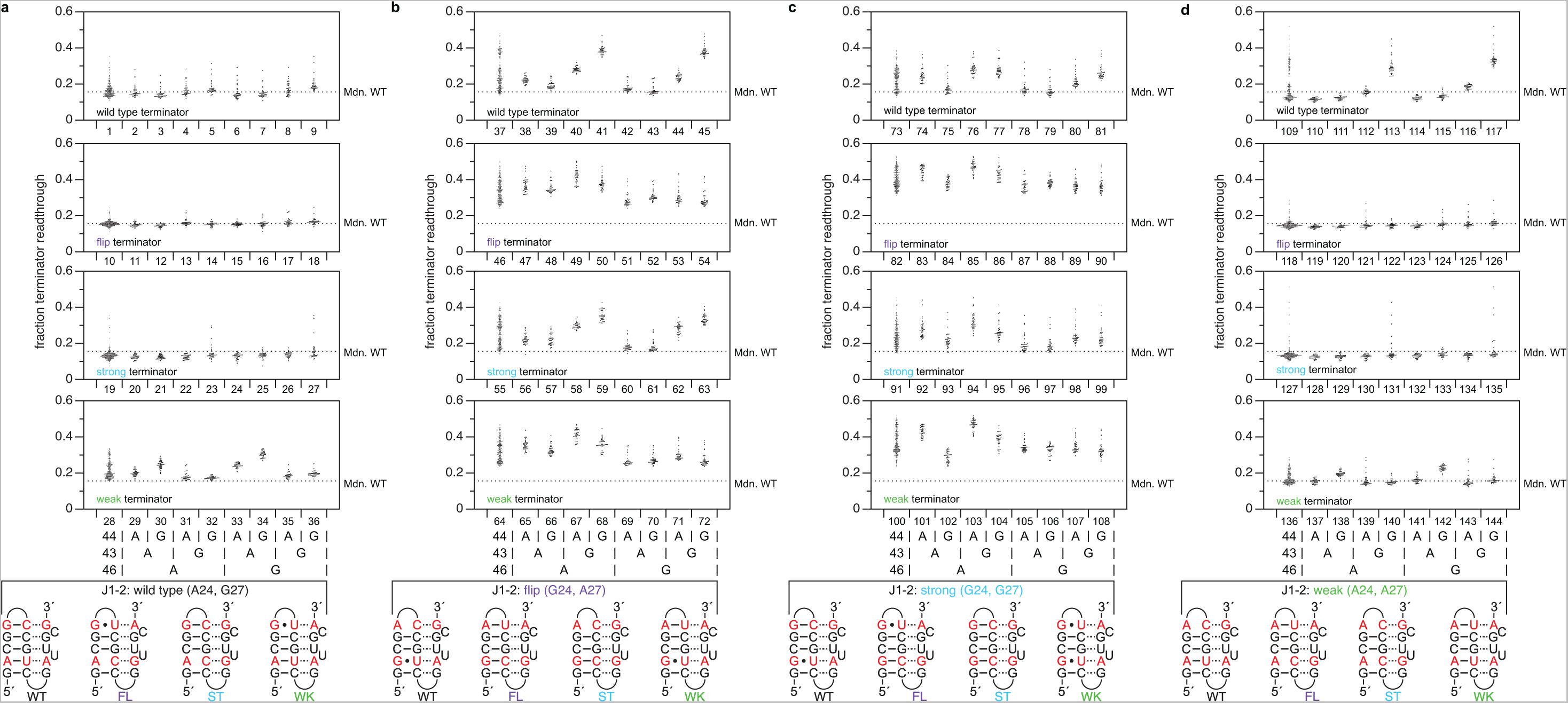
Effect of position 43, 44, and 46 variations on terminator efficiency. **a-d**, Plots showing how the composition of nucleotides 43, 44, and 46 affect fraction terminator readthrough for the wild type (a), flip (b), strong (c) and weak (d) J1-2 configurations in combination with each of the four terminators shown in Figure 3a. The median fraction terminator readthrough for variants with a WT PK and terminator is shown. All data are from the experiment shown in Figure 2. WT, wild type; FL, flip; ST, strong; WK, weak; Mdn, median.

**Supplementary Figure 4.**
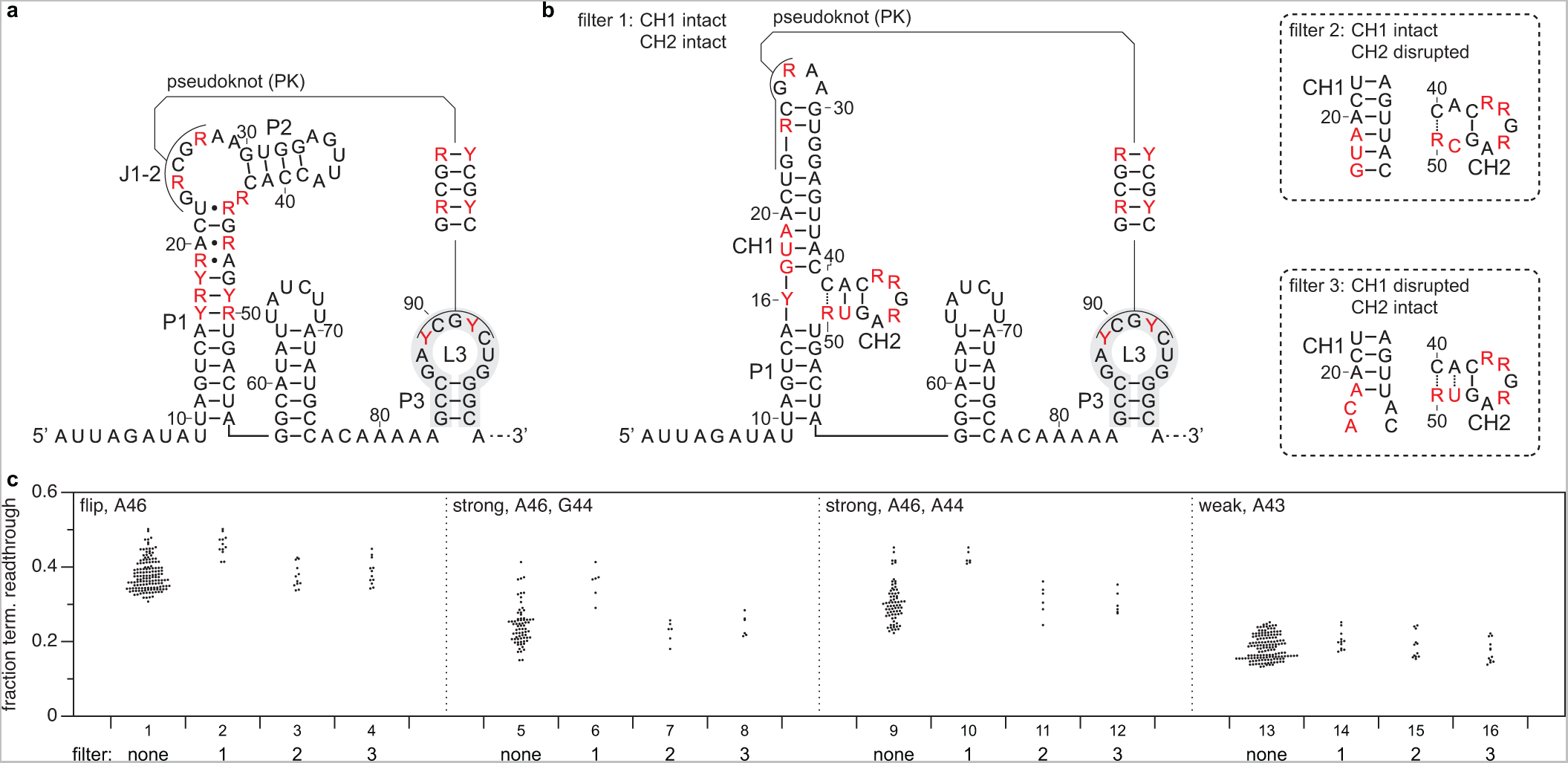
Activity of *C. beijerinckii pfl* ZTP riboswitch competitor helix variants. **a**, Sequence of the *C*. *beijerinckii pfl* ZTP riboswitch consensus variant library. **b**, Illustration of the filters that were applied in panel c. Filter 1 includes variants in which both CH1 and CH2 are intact. Filter 2 includes variants in which CH1 is intact and CH2 is disrupted. Filter 3 includes variants in which CH1 is disrupted and CH2 is intact. **c**, Plots showing the effect of CH1 and CH2 on fraction terminator readthrough in the absence of ZMP for variants in which variable nucleotides in the pseudoknot and terminator form Watson-Crick base pairs. Additional constraints on the identity of nucleotides 46, 44, and 43 are indicated. All data are from the experiment shown in Figure 2. CH1, competitor helix 1; CH2, competitor helix 2; PK, pseudoknot.

## Supplementary Tables

**Supplementary Table 1.**
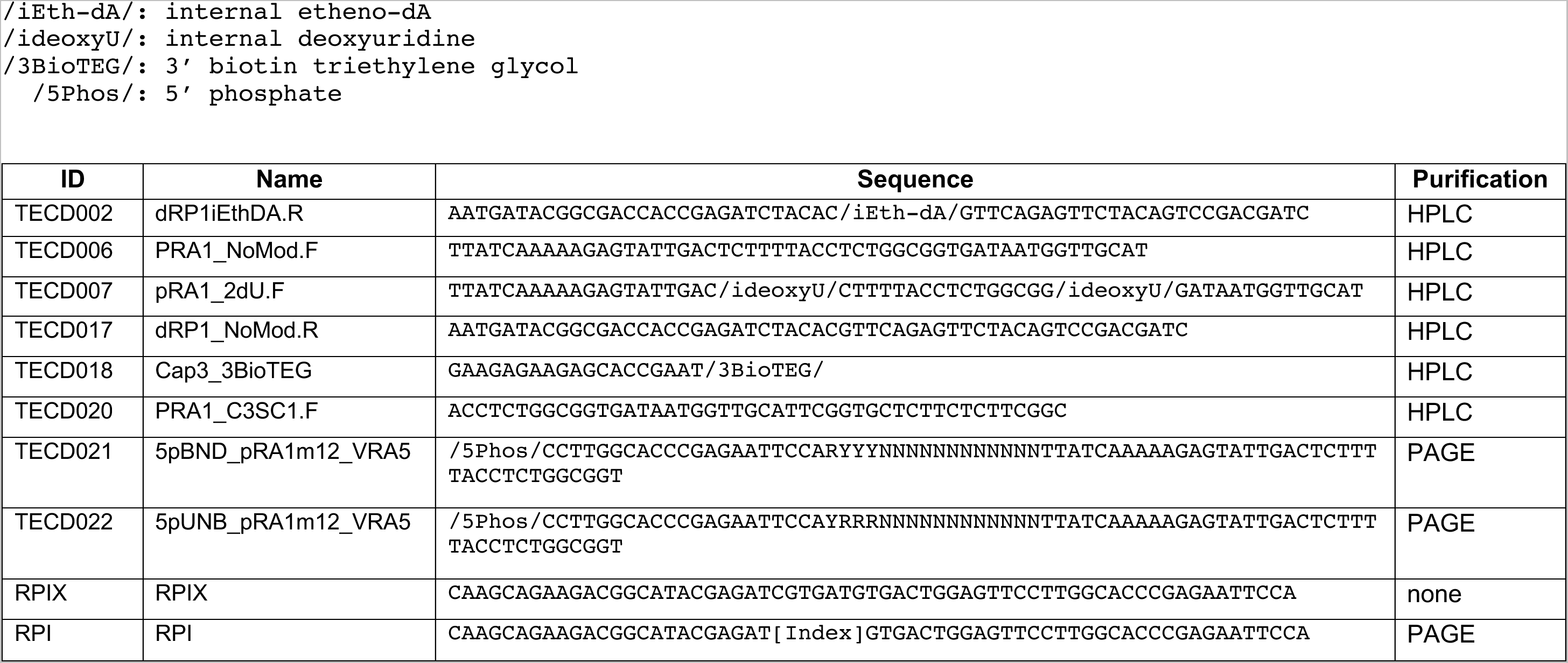
Oligonucleotides used in this study. The table below shows oligonucleotides used in this study. The modification codes presented are compatible with Integrated DNA Technology ordering.

**Supplementary Table 2.**
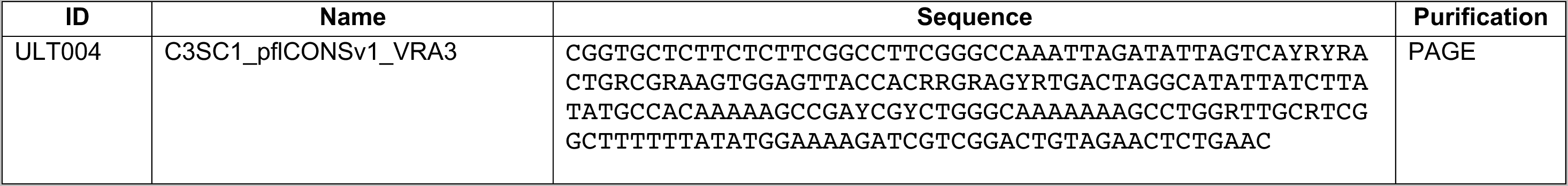
Ultramer oligonucleotide used in this study. The table below shows the sequence of the ultramer oligonucleotide used in this study.

**Supplementary Table 3.** DNA templates prepared for this study. The table below describes the DNA templates prepared for this study, including the primers and templates used, DNA modifications, whether translesion synthesis was performed, how the DNA template was purified, and the figures in which each DNA templates was used

| <u>ID</u> | <u>Fwd Primer</u> | <u>Rev Primer</u> | <u>Template</u> | <u>Modifications</u> | <u>Polymerase</u> | <u>Translesion Synthesis</u> | <u>Clean up</u> | <u>Used in Fig(s)</u> |
| --- | --- | --- | --- | --- | --- | --- | --- | --- |
| 1 | TECD006 | TECD017 | pCES002 (Kelly, Szyjka, and Strobel, 2022) | none | Vent (exo-) | N/A | Agarose gel extraction | N/A, PCR template for DNA templates 2 and 3 |
| 2 | TECD007 | TECD002 | Template 1 | -13,-30 dU;<br>Internal etheno-dA | Q5U | Yes | SPRI bead purification | 1c |
| 3 | TECD007 | TECD017 | Template 1 | -13,-30 dU | Q5U | Yes | SPRI bead purification | S1 |
| 4 | TECD020 | TECD017 | ULT004 | none | Q5 | N/A | SPRI bead purification | N/A, PCR template for DNA template 5 |
| 5 | TECD007 | TECD002 | Template 4 | -13, -30 dU;<br>Internal etheno-dA | Q5U | Yes | Polyacrylamide gel extraction, filtration, phenol /chloroform extraction | 2, 3, 4, 5, S2, S3, S4 |

**Supplementary Table 4.**
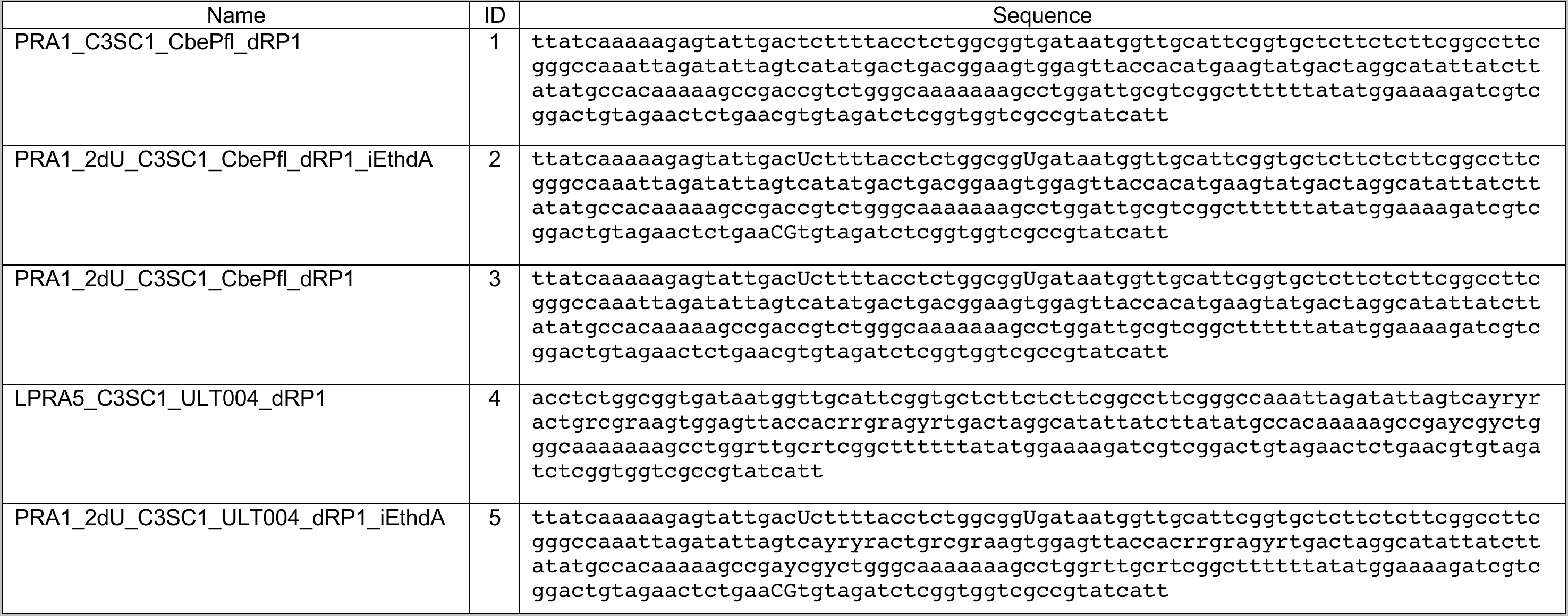
DNA template sequences. The table below contains the DNA sequences used in this study. In variants that contain an etheno-dA modification, the etheno-dA nucleotide is in the complementary (transcribed) strand between the uppercase CG.

**Supplementary Table 5.** Sequencing Read Archive (SRA) deposition table. All primary sequencing data generated in this work are freely available from the Sequencing Read Archive (http://www.ncbi.nlm.nih.gov/sra), accessible via the BioProject accession number PRJNA976257 or using the individual accession numbers below.

| SRA Accession | Sample Name | Experiment |
| --- | --- | --- |
| SAMN37456695 | ZTP_U004_0mM_ZMP_GreB_LE_Rep1 | TECdisplay data, pfl ZTP riboswitch ultramer 004, 0mM ZMP, 100 nM GreB, lambda exo treated, replicate 1 |
| SAMN37456696 | ZTP_U004_0mM_ZMP_GreB_LE_Rep2 | TECdisplay data, pfl ZTP riboswitch ultramer 004, 0mM ZMP, 100 nM GreB, lambda exo treated, replicate 2 |
| SAMN37811191 | ZTP_U004_0p031mM_ZMP_GreB_LE_Rep1 | TECdisplay data, pfl ZTP riboswitch ultramer 004, 0.0316 mM ZMP, 100 nM GreB, lambda exo treated, replicate 1 |
| SAMN37456699 | ZTP_U004_0p100mM_ZMP_GreB_LE_Rep1 | TECdisplay data, pfl ZTP riboswitch ultramer 004, 0.1 mM ZMP, 100 nM GreB, lambda exo treated, replicate 1 |
| SAMN37456700 | ZTP_U004_0p316mM_ZMP_GreB_LE_Rep1 | TECdisplay data, pfl ZTP riboswitch ultramer 004, 0.316 mM ZMP, 100 nM GreB, lambda exo treated, replicate 1 |
| SAMN37456697 | ZTP_U004_1mM_ZMP_GreB_LE_Rep1 | TECdisplay data, pfl ZTP riboswitch ultramer 004, 1mM ZMP, 100 nM GreB, lambda exo treated, replicate 1 |
| SAMN37456698 | ZTP_U004_1mM_ZMP_GreB_LE_Rep2 | TECdisplay data, pfl ZTP riboswitch ultramer 004, 1mM ZMP, 100 nM GreB, lambda exo treated, replicate 2 |
| SAMN37456701 | ZTP_U004_3p160mM_ZMP_GreB_LE_Rep1 | TECdisplay data, pfl ZTP riboswitch ultramer 004, 3.16 mM ZMP, 100 nM GreB, lambda exo treated, replicate 1 |

## Notes

### Competing Interest Statement

The authors have declared no competing interest.

## References

1 Pan, T. & Sosnick, T. RNA folding during transcription. Annu Rev Biophys Biomol Struct 35, 161–175 (2006). 10.1146/annurev.biophys.35.040405.102053

2 Lai, D., Proctor, J. R. & Meyer, I. M. On the importance of cotranscriptional RNA structure formation. RNA 19, 1461–1473 (2013). 10.1261/rna.037390.112

3 Scharfen, L. & Neugebauer, K. M. Transcription Regulation Through Nascent RNA Folding. J Mol Biol 433, 166975 (2021). 10.1016/j.jmb.2021.166975

4 Denny, S. K. & Greenleaf, W. J. Linking RNA Sequence, Structure, and Function on Massively Parallel High-Throughput Sequencers. Cold Spring Harb Perspect Biol 11 (2019). 10.1101/cshperspect.a032300

5 Marklund, E., Ke, Y. & Greenleaf, W. J. High-throughput biochemistry in RNA sequence space: predicting structure and function. Nat Rev Genet 24, 401–414 (2023). 10.1038/s41576-022-00567-5

6 Tome, J. M. et al. Comprehensive analysis of RNA-protein interactions by high-throughput sequencing-RNA affinity profiling. Nat Methods 11, 683–688 (2014). 10.1038/nmeth.2970

7 Buenrostro, J. D. et al. Quantitative analysis of RNA-protein interactions on a massively parallel array reveals biophysical and evolutionary landscapes. Nat Biotechnol 32, 562–568 (2014). 10.1038/nbt.2880

8 She, R. et al. Comprehensive and quantitative mapping of RNA-protein interactions across a transcribed eukaryotic genome. Proc Natl Acad Sci U S A 114, 3619–3624 (2017). 10.1073/pnas.1618370114

9 Denny, S. K. et al. High-Throughput Investigation of Diverse Junction Elements in RNA Tertiary Folding. Cell 174, 377–390 e320 (2018). 10.1016/j.cell.2018.05.038

10 Becker, W. R. et al. High-Throughput Analysis Reveals Rules for Target RNA Binding and Cleavage by AGO2. Mol Cell 75, 741–755 e711 (2019). 10.1016/j.molcel.2019.06.012

11 Jarmoskaite, I. et al. A Quantitative and Predictive Model for RNA Binding by Human Pumilio Proteins. Mol Cell 74, 966–981 e918 (2019). 10.1016/j.molcel.2019.04.012

12 Wu, M. J., Andreasson, J. O. L., Kladwang, W., Greenleaf, W. & Das, R. Automated Design of Diverse Stand-Alone Riboswitches. ACS Synth Biol 8, 1838–1846 (2019). 10.1021/acssynbio.9b00142

13 Yesselman, J. D. et al. Sequence-dependent RNA helix conformational preferences predictably impact tertiary structure formation. Proc Natl Acad Sci U S A 116, 16847–16855 (2019). 10.1073/pnas.1901530116

14 Andreasson, J. O. L., Savinov, A., Block, S. M. & Greenleaf, W. J. Comprehensive sequence-to-function mapping of cofactor-dependent RNA catalysis in the glmS ribozyme. Nat Commun 11, 1663 (2020). 10.1038/s41467-020-15540-1

15 Bonilla, S. L. et al. High-throughput dissection of the thermodynamic and conformational properties of a ubiquitous class of RNA tertiary contact motifs. Proc Natl Acad Sci U S A 118 (2021). 10.1073/pnas.2109085118

16 Ober-Reynolds, B. et al. High-throughput biochemical profiling reveals functional adaptation of a bacterial Argonaute. Mol Cell 82, 1329–1342 e1328 (2022). 10.1016/j.molcel.2022.02.026

17 Sadee, C. et al. A comprehensive thermodynamic model for RNA binding by the Saccharomyces cerevisiae Pumilio protein PUF4. Nat Commun 13, 4522 (2022). 10.1038/s41467-022-31968-z

18 Shin, J. H., Bonilla, S. L., Denny, S. K., Greenleaf, W. J. & Herschlag, D. Dissecting the energetic architecture within an RNA tertiary structural motif via high-throughput thermodynamic measurements. Proc Natl Acad Sci U S A 120, e2220485120 (2023). 10.1073/pnas.2220485120

19 Serganov, A. & Nudler, E. A decade of riboswitches. Cell 152, 17–24 (2013). 10.1016/j.cell.2012.12.024

20 McCown, P. J., Corbino, K. A., Stav, S., Sherlock, M. E. & Breaker, R. R. Riboswitch diversity and distribution. RNA 23, 995–1011 (2017). 10.1261/rna.061234.117

21 Hallberg, Z. F., Su, Y., Kitto, R. Z. & Hammond, M. C. Engineering and In Vivo Applications of Riboswitches. Annu Rev Biochem 86, 515–539 (2017). 10.1146/annurev-biochem-060815-014628

22 Porter, E. B., Polaski, J. T., Morck, M. M. & Batey, R. T. Recurrent RNA motifs as scaffolds for genetically encodable small-molecule biosensors. Nat Chem Biol 13, 295–301 (2017). 10.1038/nchembio.2278

23 Topp, S. & Gallivan, J. P. Emerging applications of riboswitches in chemical biology. ACS Chem Biol 5, 139–148 (2010). 10.1021/cb900278x

24 Braselmann, E. et al. A multicolor riboswitch-based platform for imaging of RNA in live mammalian cells. Nat Chem Biol 14, 964–971 (2018). 10.1038/s41589-018-0103-7

25 Kim, P. B., Nelson, J. W. & Breaker, R. R. An ancient riboswitch class in bacteria regulates purine biosynthesis and one-carbon metabolism. Mol Cell 57, 317–328 (2015). 10.1016/j.molcel.2015.01.001

26 Tran, B. et al. Parallel Discovery Strategies Provide a Basis for Riboswitch Ligand Design. Cell Chem Biol 27, 1241–1249 e1244 (2020). 10.1016/j.chembiol.2020.07.021

27 Perkins, K. R., Atilho, R. M., Moon, M. H. & Breaker, R. R. Employing a ZTP Riboswitch to Detect Bacterial Folate Biosynthesis Inhibitors in a Small Molecule High-Throughput Screen. ACS Chem Biol 14, 2841–2850 (2019). 10.1021/acschembio.9b00713

28 Kelly, S. L., Szyjka, C. E. & Strobel, E. J. Purification of synchronized Escherichia coli transcription elongation complexes by reversible immobilization on magnetic beads. J Biol Chem 298, 101789 (2022). 10.1016/j.jbc.2022.101789

29 Strobel, E. J., Lis, J. T. & Lucks, J. B. Chemical roadblocking of DNA transcription for nascent RNA display. J Biol Chem 295, 6401–6412 (2020). 10.1074/jbc.RA120.012641

30 Pupov, D., Ignatov, A., Agapov, A. & Kulbachinskiy, A. Distinct effects of DNA lesions on RNA synthesis by Escherichia coli RNA polymerase. Biochem Biophys Res Commun 510, 122–127 (2019). 10.1016/j.bbrc.2019.01.062

31 Strobel, E. J. Efficient Linear dsDNA Tagging Using Deoxyuridine Excision*. Chembiochem 22, 3214–3224 (2021). 10.1002/cbic.202100425

32 Alon, S. et al. Barcoding bias in high-throughput multiplex sequencing of miRNA. Genome Res 21, 1506–1511 (2011). 10.1101/gr.121715.111

33 Hafner, M. et al. RNA-ligase-dependent biases in miRNA representation in deep-sequenced small RNA cDNA libraries. RNA 17, 1697–1712 (2011). 10.1261/rna.2799511

34 Smola, M. J., Rice, G. M., Busan, S., Siegfried, N. A. & Weeks, K. M. Selective 2’-hydroxyl acylation analyzed by primer extension and mutational profiling (SHAPE-MaP) for direct, versatile and accurate RNA structure analysis. Nat Protoc 10, 1643–1669 (2015). 10.1038/nprot.2015.103

35 Spitale, R. C. et al. Structural imprints in vivo decode RNA regulatory mechanisms. Nature 519, 486–490 (2015). 10.1038/nature14263

36 Strobel, E. J., Watters, K. E., Nedialkov, Y., Artsimovitch, I. & Lucks, J. B. Distributed biotin-streptavidin transcription roadblocks for mapping cotranscriptional RNA folding. Nucleic Acids Res 45, e109 (2017). 10.1093/nar/gkx233

37 Lucas, J. K., Gruenke, P. R. & Burke, D. H. Minimizing amplification bias during reverse transcription for in vitro selections. RNA 29, 1301–1315 (2023). 10.1261/rna.079650.123

38 Szyjka, C. E. & Strobel, E. J. Observation of coordinated RNA folding events by systematic cotranscriptional RNA structure probing. Nat Commun 14, 7839 (2023). 10.1038/s41467-023-43395-9

39 Verwilt, J., Mestdagh, P. & Vandesompele, J. Artifacts and biases of the reverse transcription reaction in RNA sequencing. RNA 29, 889–897 (2023). 10.1261/rna.079623.123

40 Strobel, E. J., Cheng, L., Berman, K. E., Carlson, P. D. & Lucks, J. B. A ligand-gated strand displacement mechanism for ZTP riboswitch transcription control. Nat Chem Biol 15, 1067–1076 (2019). 10.1038/s41589-019-0382-7

41 Reuter, J. S. & Mathews, D. H. RNAstructure: software for RNA secondary structure prediction and analysis. BMC Bioinformatics 11, 129 (2010). 10.1186/1471-2105-11-129

42 Trausch, J. J., Marcano-Velazquez, J. G., Matyjasik, M. M. & Batey, R. T. Metal Ion-Mediated Nucleobase Recognition by the ZTP Riboswitch. Chem Biol 22, 829–837 (2015). 10.1016/j.chembiol.2015.06.007

43 Nadon, J. F. et al. Site-specific photolabile roadblocks for the study of transcription elongation in biologically complex systems. Commun Biol 5, 457 (2022). 10.1038/s42003-022-03382-0

44 Strobel, E. J. Preparation of E. coli RNA polymerase transcription elongation complexes by selective photoelution from magnetic beads. J Biol Chem 297, 100812 (2021). 10.1016/j.jbc.2021.100812

45 Strobel, E. J. Isolation of E. coli RNA polymerase transcription elongation complexes by selective solid-phase photoreversible immobilization. Methods Enzymol 691, 223–250 (2023). 10.1016/bs.mie.2023.03.019

46 Perederina, A. A. et al. Cloning, expression, purification, crystallization and initial crystallographic analysis of transcription elongation factors GreB from Escherichia coli and Gfh1 from Thermus thermophilus. Acta Crystallogr Sect F Struct Biol Cryst Commun 62, 44–46 (2006). 10.1107/S1744309105040297

47 Torgerson, C. D., Hiller, D. A., Stav, S. & Strobel, S. A. Gene regulation by a glycine riboswitch singlet uses a finely tuned energetic landscape for helical switching. RNA 24, 1813–1827 (2018). 10.1261/rna.067884.118

48 Torgerson, C. D., Hiller, D. A. & Strobel, S. A. The asymmetry and cooperativity of tandem glycine riboswitch aptamers. RNA 26, 564–580 (2020). 10.1261/rna.073577.119

49 Focht, C. M. & Strobel, S. A. Efficient quantitative monitoring of translational initiation by RelE cleavage. Nucleic Acids Res 50, e105 (2022). 10.1093/nar/gkac614

50 Focht, C. M., Hiller, D. A., Grunseich, S. G. & Strobel, S. A. Translation regulation by a guanidine-II riboswitch is highly tunable in sensitivity, dynamic range, and apparent cooperativity. RNA 29, 1126–1139 (2023). 10.1261/rna.079560.122

51 Thompson, J. R., Marcelino, L. A. & Polz, M. F. Heteroduplexes in mixed-template amplifications: formation, consequence and elimination by ‘reconditioning PCR’. Nucleic Acids Res 30, 2083–2088 (2002). 10.1093/nar/30.9.2083

52 Strobel, E. J., Kelly, S. L. & Szyjka, C. E. Isolation of synchronized E. coli elongation complexes for solid-phase and solution-based in vitro transcription assays. Methods Enzymol 675, 159–192 (2022). 10.1016/bs.mie.2022.07.008

53 Borukhov, S., Sagitov, V. & Goldfarb, A. Transcript cleavage factors from E. coli. Cell 72, 459–466 (1993). 10.1016/0092-8674(93)90121-6

54 Komissarova, N. & Kashlev, M. Transcriptional arrest: Escherichia coli RNA polymerase translocates backward, leaving the 3’ end of the RNA intact and extruded. Proc Natl Acad Sci U S A 94, 1755–1760 (1997). 10.1073/pnas.94.5.1755

55 Nudler, E., Mustaev, A., Lukhtanov, E. & Goldfarb, A. The RNA-DNA hybrid maintains the register of transcription by preventing backtracking of RNA polymerase. Cell 89, 33–41 (1997). 10.1016/s0092-8674(00)80180-4

56 Laptenko, O., Lee, J., Lomakin, I. & Borukhov, S. Transcript cleavage factors GreA and GreB act as transient catalytic components of RNA polymerase. EMBO J 22, 6322–6334 (2003). 10.1093/emboj/cdg610

57 Artsimovitch, I. & Landick, R. Pausing by bacterial RNA polymerase is mediated by mechanistically distinct classes of signals. Proc Natl Acad Sci U S A 97, 7090–7095 (2000). 10.1073/pnas.97.13.7090

58 Larson, M. H. et al. A pause sequence enriched at translation start sites drives transcription dynamics in vivo. Science 344, 1042–1047 (2014). 10.1126/science.1251871

59 Vvedenskaya, I. O. et al. Interactions between RNA polymerase and the “core recognition element” counteract pausing. Science 344, 1285–1289 (2014). 10.1126/science.1253458

60 Meyer, I. M. & Miklos, I. Co-transcriptional folding is encoded within RNA genes. BMC Mol Biol 5, 10 (2004). 10.1186/1471-2199-5-10

61 Roberts, R. W. & Szostak, J. W. RNA-peptide fusions for the in vitro selection of peptides and proteins. Proc Natl Acad Sci U S A 94, 12297–12302 (1997). 10.1073/pnas.94.23.12297

62 Hanes, J. & Pluckthun, A. In vitro selection and evolution of functional proteins by using ribosome display. Proc Natl Acad Sci U S A 94, 4937–4942 (1997). 10.1073/pnas.94.10.4937

63 MacPherson, I. S., Temme, J. S. & Krauss, I. J. DNA display of folded RNA libraries enabling RNA-SELEX without reverse transcription. Chem Commun (Camb) 53, 2878–2881 (2017). 10.1039/c6cc09991b

64 Mohsen, M. G., Midy, M. K., Balaji, A. & Breaker, R. R. Exploiting natural riboswitches for aptamer engineering and validation. Nucleic Acids Res 51, 966–981 (2023). 10.1093/nar/gkac1218

65 Strobel, E. J. Preparation and Characterization of Internally Modified DNA Templates for Chemical Transcription Roadblocking. Bio Protoc 11, e4141 (2021). 10.21769/BioProtoc.4141

66 Jolivet, P. & Foley, J. SPRI bead mix. *protocols.io* (2020). dx.doi.org/10.17504/protocols.io.bnz4mf8w

67 Mahat, D. B. et al. Base-pair-resolution genome-wide mapping of active RNA polymerases using precision nuclear run-on (PRO-seq). Nat Protoc 11, 1455–1476 (2016). 10.1038/nprot.2016.086

68 Chen, S., Zhou, Y., Chen, Y. & Gu, J. fastp: an ultra-fast all-in-one FASTQ preprocessor. Bioinformatics 34, i884–i890 (2018). 10.1093/bioinformatics/bty560

69 Ren, A., Rajashankar, K. R. & Patel, D. J. Global RNA Fold and Molecular Recognition for a pfl Riboswitch Bound to ZMP, a Master Regulator of One-Carbon Metabolism. Structure 23, 1375–1381 (2015). 10.1016/j.str.2015.05.016

70 Jones, C. P. & Ferre-D’Amare, A. R. Recognition of the bacterial alarmone ZMP through long-distance association of two RNA subdomains. Nat Struct Mol Biol 22, 679–685 (2015). 10.1038/nsmb.3073

71 Pettersen, E. F. et al. UCSF Chimera--a visualization system for exploratory research and analysis. J Comput Chem 25, 1605–1612 (2004). 10.1002/jcc.20084

